# Tuning of granulopoietic signaling by *de novo* designed agonists

**DOI:** 10.1101/2023.11.25.568662

**Authors:** Timo Ullrich, Christoph Pollmann, Malte Ritter, Jérémy Haaf, Narges Aghaallaei, Ivan Tesakov, Maya El-Riz, Kateryna Maksymenko, Valeriia Hatskovska, Sergey Kandabarau, Maksim Klimiankou, Claudia Lengerke, Karl Welte, Birte Hernandez-Alvarez, Patrick Müller, Andrei Lupas, Jacob Piehler, Julia Skokowa, Mohammad ElGamacy

## Abstract

Enhancing cytokine-based therapies by systematically tuning how an agonist associates its receptor is emerging as a powerful new concept in drug discovery. Here, we report the design and characterization of agonists that tune the granulocyte-colony stimulating factor receptor (G-CSFR) activity, which is central for the proliferation and granulocytic differentiation of hematopoietic stem cells. Using design agonists, we study the impact of varying the receptor-binding affinity and dimerization geometry on receptor association, downstream signaling, and cellular response. Hence, we achieved agonists with altered signaling specificities that are hyper-thermostable, can outcompete the native ligand (G-CSF), and bias granulopoietic differentiation over triggering proliferation. Furthermore, the design agonists differentially modulate the kinetics and amplitudes of signal transduction pathways, and gene expression patterns. Unlike G-CSF, they achieve selective activation of gene sets with hematopoietic functions with minimal unwanted effects on immunomodulatory signaling. These findings demonstrate the potential of dissecting the complex G-CSFR signaling, and open up ways for new therapeutic applications for designed cytokines.

**Graphical abstract:** 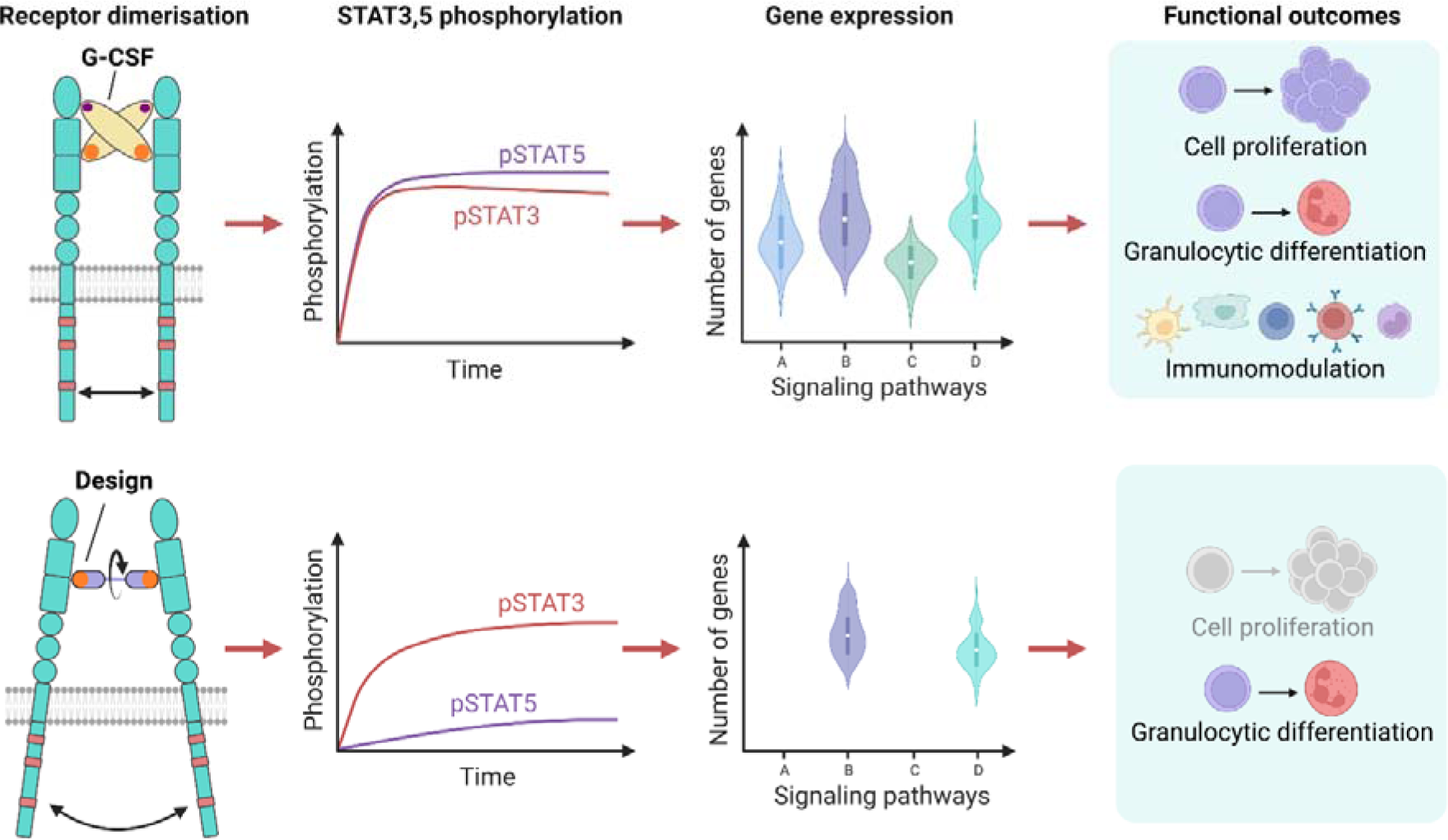

## Introduction

Cytokine receptor activation typically triggers several signaling pathways that results in different cellular responses depending on target cells [1, 2]. Synthetic ligands that engage cytokine receptors in non-native ways can selectively bias the signaling outcomes, overcoming the functional pleiotropy of native agonists [3]. Modulation of class I/II cytokine receptors is of considerable interest for therapeutic intervention, but is hampered by the often broad and pleiotropic responses [4–8]. Decomposing these signaling outcomes through synthetic agonists can thus unlock previously untapped therapeutic applications [9].

The observed functional plasticity of natural ligands [10] has inspired efforts to systematically bias signaling specificity using design agonists purposefully engineered to possess optimal pharmaceutical properties [9, 11]. For instance, affinity-engineering of IL-2 [12] or IL-22 [13] variants possessed immunomodulatory activity or altered tissue specificity when compared to the respective natural ligands. This highlight the effect of the ligand’s receptor-binding affinity on downstream signaling [14, 15]. In addition to affinity, the receptors’ association geometry can provide another layer of control over downstream signaling [16–19], which was demonstrated through synthetic ligands. Specifically, diabodies and DARPins dimerizing the erythropoietin receptor (EPOR) in non-native orientations could induce a range of differential signaling outcomes [20, 21]. Alternatively, using antibody fragment fusions, signal tuning was demonstrated for heterodimeric receptor complexes of the IL-2 receptor (IL-2Rβ:γ_c_) and the type I interferon receptor (IFNAR1:IFNAR2) [22].

In this study, we explore the tunability of granulocyte-colony stimulating factor receptor (G-CSFR). Under physiological conditions, the native ligand (G-CSF) homodimerizes G-CSFR to induce signaling, which is crucial for granulopoiesis, and the regulation of hematopoietic stem and progenitor cells (HSPCs), and immune cells [23–30]. Despite the clinical adoption of recombinant G-CSF (rhG-CSF) for treating neutropenia and stem cell mobilization [31, 32], its therapeutic applications are still limited due to its capacity to inadvertently activate, e.g., different types of immune cells, tumor cells, osteoclasts, which express G-CSFR. By designing G-CSFR agonists that associate the receptor subunits at varying affinities and geometries we were able to control the downstream phosphorylation of key signal transducers and activators of transcription (STATs). Based on characterizing the biophysical properties of our design agonists and investigating their impact on receptor dimerization in the plasma membrane, in conjunction with intracellular signal activation, and cellular activation *in vitro* and *in vivo*, we highlight the potential to tune G-CSFR signaling towards differentiation, while avoiding its unwanted non-hematopoietic effects.

## Material and methods

### Computationally-guided library generation for affinity maturation

Two libraries of mutants of 6 amino acid positions on the Boskar4 receptor-binding sites were generated using either site-saturation mutagenesis or computational design. The mutated positions on the Boskar4 surface were chosen based on two criteria (Fig. S1A). The positions in Boskar4 were determined which had the smallest average distance to G-CSFR in a modeled complex between the Boskar4 ensemble (PDB: 7NY0, [33]) and G-CSFR (PBD:2D9Q, [29]). Residues known to be critical for G-CSF activity were excluded from the final choice [34]. For the saturation library, high diversity degenerated codons were chosen which contained the original amino acid and other random amino acids constructing a library of a theoretical diversity (number of distinct gene variants) of 4.0×10^6^ and an amino-acid diversity (number of distinct protein variants) of 3.2×10^6^. Finally, the library was generated by PCR (Table S2 and S3) using one pair of primers (primer 3 and 4), each containing 3 degenerated sites to linearize an *E. coli* display system plasmid [35] encoding a fusion between the N-terminal part of Intimin and Boskar4 (pNB4, Table S4, Fig. S1B, C). The obtained PCR product was purified with the Wizard® SV Gel and PCR Clean-Up System (A928, Promega) and blunt end ligation was performed followed by electroporation of freshly made competent cells [36] to yield 3.6×10^7^ total transformants. The *E. coli* strain DH10B T1^R^ (C640003, Invitrogen) was used for all display experiments.

In more detail, prior to transformation, 0.4 U/µL T4 polynucleotide kinase (EK0031, Thermo Scientific™) was incubated with 5 ng/µL of linearized PCR product for 30 min at 37 °C, according to manufacturer instructions. After that, 0.25 U/µL T4 DNA ligase (EL0011, Thermo Scientific™) was added and incubated overnight at 16 °C. The ligation product was purified with the same PCR Clean-Up System mentioned above and eluted into 20 µL filtered ddH_2_O. The complete 20 µL were transformed into electrocompetent *E. coli* as described in Tu *et al.* with the following changes [36]. In short, 50 mL of freshly grown *E. coli* exhibiting an optical density at 600 nm (*OD_600_*) of ∼ 0.6 were washed two times with 35 mL filtered H_2_O. After washing, the bacterial pellet was resuspended with the purified ligation mix. The electroporation was performed at 1250 V and 5 ms. The cells were grown in 1 mL SOC Outgrowth Medium (B9020, NEB) for 2 h in a standard glass tube at 30 °C with 160 rpm. A 10-fold dilution series of the cells was made in SOC-medium, and they were plated on agar plates containing 34 µg/mL chloramphenicol and 2 % (w/v) D-glucose to estimate the total number of transformants. The rest of the sample was plated on five agar plates of the same type and incubated at 30 °C for about 20 h. The bacterial lawn was scraped from the plates and well mixed in 10 mL lysogeny broth (LB) medium. From 500 µL of bacterial suspension, the display plasmid containing the Boskar4 library was isolated and analyzed by Sanger sequencing with primers 19 and 20.

For the designed library, Damietta spl application [37] was used to estimate the free energy change of each single mutants of the same positions and same amino acids per position as the control library in a non-combinatorial manner. For each of the six positions, five amino acids with the lowest free energy change plus the original amino acid of Boskar4 at the corresponding position were used to define the designed, size-reduced library with a theoretical diversity of 3.89×10^4^. The library was generated by PCR the same way as the original one but using three pairs of primers (primer 5 to 10, Table S2) such that the final library contained 15.6% of desired amino acids. The primer composition was designed by using the online tool of SwiftLib [38]. The designed library had a final theoretical diversity of 2.5×10^5^, an amino-acid diversity of 2.3×10^5^and 2.7×10^7^total transformants were yielded.

### Screening procedure for enhanced binders

The FACS-based screening of bacterially-displayed Boskar4 variants followed a slightly adapted procedure from that described by Salema *et al.* [35]. In brief, G-CSFR was first fluorescently labelled. This was done by incubating 100 µg of recombinant human G-CSFR (381-GR/CF, R&D Systems) in 950 µL PBS for 2 h at room temperature (RT) with Biotin-NHS (H1759, Sigma) in a 1:20 molar ratio (G-CSFR:Biotin-NHS). The reaction was stopped with 50 µL Tris (1 M pH 7.5). After 1 h of incubation on ice, the product was purified with a desalting column (Sephadex G25 PD-10, GE Healthcare), and the elution fractions were concentrated with Amicon Ultra centrifugal filter units (3 kDa, UFC800324, Merck) to a concentration of 0.24 mg/mL (3.6 µM). Aliquots of the biotinylated protein (herein referred to as BioGCSFR) were stored at -20 °C until use.

For the first sorting, DH10B T1^R^ *E. coli* cells were transformed with the display-vector pN containing a Boskar4 library. The bacteria were freshly grown on LB agar plates containing 34 µg/mL chloramphenicol and 2% (w/v) D-glucose at 30 °C yielding ∼3.0×10^8^ total transformants. Following the harvesting of bacteria from the agar plates, 1 mL of resuspended bacterial culture exhibiting an *OD_600_*of 3.3 was subjected to two successive washes, each involving 1 mL of LB medium. Subsequently, the culture was diluted to an initial *OD_600_* of 0.33 by adding it to 10 mL of LB medium containing 34 µg/mL of chloramphenicol. The cells were cultivated at a temperature of 30 °C and an agitation rate of 160 rpm. After a duration of 1 h, 50 µM of isopropyl β-D-1-thiogalactopyranoside (IPTG) was introduced into the culture, and the cultivation was continued for an additional 3 h under the same growth conditions. Cells equivalent to 1 mL of a cell resuspension possessing an *OD_600_* of 1 were collected, subjected to two washes with 1000 µL of phosphate-buffered saline (PBS), and eventually resuspended in 400 µL of PBS. 190 µL of this cell suspension was incubated with 10 nM BioGCSFR for 1 h at room temperature, followed by one wash with 1000 µL PBS. The washed cells were resuspended in 200 µL PBS containing 0.375 µL PE-Streptavidin (405203, BioLegend) and were subsequently incubated for 30 min at 4 °C. Finally, the sample was washed one more time with 1000 µL PBS and resuspended in 1000 µL PBS. FACS was performed with a BD FACSMelody™ Cell Sorter having set the PTM voltage for PE to 517 V, SSC to 426 V and a SSC threshold to 673 V. No gating was applied. All flow cytometry results were analyzed using FlowJo™ v10.1 Software (BD Life Sciences). 300 µL of the sample was mixed with 4 mL ice cold PBS and the flow rate was adjusted to end up with an event rate of ∼8000 events/sec. 100,000 events were recorded per sample. The top ∼0.1 % of the total population depending on a SSC-A/PE-A plot were sorted into a 1.5 mL tube provided with 200 µL LB medium and cooled to 5 °C. The sorting was continued until at least 20-fold more processed events were screened than the theoretical diversity of the correspondent library was. All sorted cells were plated on appropriate LB agar plates and incubated for about 20 h at 30 °C. Those plates were harvested and grown over night in LB medium containing 34 µg/mL chloramphenicol and 2% (w/v) D-glucose at 30 °C under static conditions. From this culture, 1 mL of resuspended bacteria with an *OD_600_*of 3.3 used for the next selection cycle performed in the same way as the first one.

This was followed by evaluating the relative binding of the top clones using plate-based assays. At this stage, single colonies of bacteria pools which were enriched for 3 or 5 cycles of FACS-based sorting (as described in the section above) were grown overnight in 100 µL LB (supplemented with 34 µg/mL chloramphenicol and 50 µM IPTG) in a 96-well plate sealed with Breathe-Easy ® sealing membrane (Z380059, Sigma-Aldrich) at 30 °C with 1100 rpm on a table top shaker (Thermomixer comfort, Eppendorf) covered with aluminum foil. On the next day, the cells were centrifuged at 3200 g for 5 min, the supernatant was decanted, and the cells were resuspended in 150 µL PBS. The washing was repeated the same way one more time, and the cells were resuspended in 50 µL PBS containing 10 nM BioGCSFR. After the plate was incubated for 1 h at RT, the cells were washed once with 200 µL PBS and resuspended in 50 µL PBS. To visualize the binding activity of each displayed Boskar4 variant to BioGCSFR, two separate secondary stainings, one fluorescent and one chemiluminescent, were performed as follows:

For the fluorescent staining, 25 µL of the cells were mixed with 25 µL PBS containing 0.094 µL PE-streptavidin followed by a 30 min incubation at 4 °C in the dark. After washing the cells one more time with PBS, the *OD_600_* and the fluorescence (excitation wavelength = 495 nm, emission wavelength = 574 nm) were measured on a plate reader (Synergy H4 Hybrid Microplate Reader, BioTek).

For the chemiluminescence stain, 25 µL of the cells were mixed with 25 µL PBS containing 1.25 µg/mL Avidin-HRP conjugate (Invitrogen, 434423) followed by a 30 min incubation at 4 °C in the dark. Afterwards the cells were washed once with 200 µL PBS and resuspended in 100 µL PBS. The *OD_600_* of each well was measured with the plate reader. In a separate dark plate (655097, Greiner Bio One) 12.5 µL cell suspension was added to 75 µL PBS. Then 12.5 µL ECL-substrate-solution (1705061, Bio-Rad) was pipetted quickly into each well, the plate was sealed with parafilm and mixed vigorously for 10 seconds on a vortex orbital shaker. Then the luminescence of each well was measured on the plate reader (Synergy H4 Hybrid Microplate Reader, Agilent BioTek) set to an integration time of 1 second, a gain of 135, normal read speed and a 100-millisecond delay, and a read height of 1 mm.

Both readouts were analyzed by normalizing to the corresponding cell density and calculating a z-score for each sample over all wells of the same condition. From the top 5 variants of each condition, plasmids were isolated with a plasmid preparation kit (740588, Macherey-Nagel) from a 3 mL overnight culture grown at 30 °C with 160 rpm in LB with 34 µg/mL chloramphenicol and 2% (w/v) D-glucose.

Finally, to obtain the expression-normalized binding of the top clones with unique sequences another FACS experiment was carried out. The individual bacteria clones carrying the display vector pN with Intimin fused either to Boskar4 or enhanced Boskar4 variant (bv1 to bv16) were grown over night at 30 °C with 160 rpm in 3 mL LB with the addition of 34 µg/mL chloramphenicol and 50 µM IPTG. The cells were pelleted at 4000 g for 3 min, the supernatant was discarded and the sample was resuspended in 1000 µL filtered PBS. This washing step was repeated one more time and the cells were resuspended in 1000 µL PBS. 45 µL of cells suspension were incubated for 90 min at RT in a total volume of 55 µL with 10 nM BioGCSFR and 1:200 Myc-Tag (9B11) Mouse mAb (2276, Cell Signaling Technology). After one wash with 500 µL PBS, the cells were resuspended in 200 μL of PBS containing 0.375 µL PE-Streptavidin (405203, BioLegend) and 0.5 μL of anti-mouse-IgG1 conjugated to Alexa 488 Fluor (A-21202, Invitrogen) and incubated for 30 min at 4 °C in the dark. Finally, the sample was washed one more time with 500 µL PBS, resuspended in 1000 µL PBS, and 100000 events were measured by FACS.

### Protein expression and purification

First, a set of affinity-enhanced Boskar4 (B4) variant genes (bv1, bv2, bv6, bv8, bv15, and to bv16), which were identified from the affinity maturation screens, were used to generate short tandem fusions (st2, compare Fig. S4). In more detail, a single Boskar4 variant gene was amplified by PCR from pN separately using two different primer pairs (15 and 16, or 17 and 18; Table S2). The two obtained PCR fragments were assembled by NEBuilder® HiFi DNA Assembly Master Mix (E2621, New England BioLabs) into a pET28a backbone generated by PCR with primers 13 and 14. Additionally, the monomer of bv6 was cloned from pN to pET28a.

*E. coli* BL21(DE3) carrying a certain expression construct were grown to an *OD_600_* of ∼0.6 in LB medium containing 40 µg/mL kanamycin at 37 °C and 160 rpm. Cells were induced with 0.5 mM IPTG and the protein expression was performed at 25 °C and 160 rpm for about 16 h. The cells were harvested at 8000 g for 30 min, the supernatant was decanted, and the pellet was lysed by sonication in lysis buffer (50 mM Tris pH8, 100 mM (for st2 designs) or 1 M NaCl (for ori designs), 20 µg/mL DNase (#A3778, ITW Reagents) and protease inhibitor cocktail (04693132001, Roche)). The lysate was centrifuged at 16000 g for 50 min, and the supernatant was used to perform Nickel Immobilized Metal Affinity Chromatography (IMAC). Finally, size exclusion chromatography (SEC) was performed with PBS on the concentrated sample with a HiLoad 16/600, Superdex 200 pg (GE28-9893-35, Merck) column or other if mentioned in the corresponding legend description. Fractions that were supposed to contain the protein of the expected size were concentrated by ultrafiltration (10 kDa, UFC9010, Millipore), and aliquots were frozen at -20 °C until use.

### Protein folding and thermal stability

Circular dichroism (CD) spectra were recorded using a JASCO J-810 spectrometer with 0.3 mL samples at a protein concentration of 0.1 mg/mL in PBS buffer (pH 7.1) placed in 2 mm path length cuvettes. The spectral scans of the mean residual ellipticity of three accumulations were measured at a resolution of at least 0.5 nm over a range of 250-190 nm. Nano differential scanning fluorimetry (nanoDSF) was conducted on a Prometheus NT.48 and standard Prometheus capillaries (Nanotemper, PR-C002). A temperature ramp of 1 °C/min from 20 °C to 110 °C to 20 °C was applied for melting and cooling, using 1 mg/mL protein samples in PBS (unless otherwise specified in the corresponding figure legend).

### Affinity determination with surface plasmon resonance

To determine the receptor-binding affinity of Boskar4 variants, multi-cycle kinetics experiments were performed on a Biacore X100 system (GE Healthcare Life Sciences). Recombinant human G-CSFR (381-GR/CF, R&D Systems) was diluted to 50 μg/mL in 10 mM acetate buffer at pH 5.0 and immobilized on the surface of a CM5 sensor chip (GE Healthcare 29149604) using amine coupling chemistry. The protein samples were diluted in a running buffer (PBS with 0.05% v/v Tween-20). The measurements were performed at 25°C at a flow rate of 30 μL/min. Four sequential concentrations of the sample solution were used as follows: for Boskar4; 5000 nM, 2500 nM, 625 nM and 156.3 nM, for bv6; 125 nM, 63.5 nM, 31.3 nM, 15.6 nM; for all st2 variants (bv1_st2 to bv16_st2) and oris (ori0 to ori4); 20 nM, 10 nM, 5 nM and 2.5 nM). These samples were injected over the functionalized sensor chip surface for 180 s, followed by a 600 s dissociation phase with running buffer. At the end of each run, the sensor surface was regenerated with a 60 s injection of 50 mM NaOH. The reference responses and zero-concentration sensograms were subtracted from each dataset (double-referencing). Association rate constant (*k_a_*), dissociation rate constant (*k_d_*), and apparent equilibrium dissociation constants (*K_D_*) were obtained using the Biacore X100 Evaluation Software following a 1:1 binding kinetic.

### Generation of orientation-rigging designs

To generate preliminary constructs of the orientation-rigging designs (oris) containing a rigid helix-linker fusing two binding protein monomers, two bv6 monomers were connected N-to C-terminally (helix 4 of monomer 1 to helix 1 of monomer 2) with variable lengths of poly-alanine-stretches. The obtained sequences were modeled by AF2, and four constructs (ori1, ori2, ori3 and ori4) with an increasing number of connecting alanines were settled as a starting set of designs. The obtained AF2 structures were used as input to optimize the sequence of the helix-linker region with Damietta. Depending on the construct, up to 14 positions (compare Supplementary Methods) were mutated utilizing the few-to-many-to-few combinatorial sampler of Damietta (v0.32, [37]). Analysis of the designs’ structural stability was done using a tempering molecular dynamics routine, and candidates ranked by their conformational homogeneity score, as previously described in [33]. This molecular dynamics analysis was performed for all unique variants that were generated by Damietta and the most stable variant of each construct was considered as the final design. To generate a binding protein in the same fashion that minimizes the distance of the two transmembrane domains of the bound G-CSFRs, hypothetical complexes of rigid tandem-fusions with varying poly-alanine-helix-stretches and two bound G-CSFRs were created manually. Based on the expected distance, possible steric hindrances and overall length of the rigid helix-linker, ori0 was selected and a final design was obtained as described above.

The ori expression constructs were generated the same way as explained for the short tandem fusions (compare Material and Methods section “Protein expression and purification”). In more detail, the two fragments (fr1 and fr2) encoding each one bv6 monomer and the corresponding rigid helix-linker were amplified and assembled together with the pET28a-backbone. The first fragment was generated with primer 15 plus the corresponding primer for each construct (primers 21, 23, 25, 27 or 29) and the second fragment with primer 18 plus the corresponding primer (primers 22, 24, 26, 28 or 30, Table S2).

### NFS-60 cell proliferation assays

NFS-60 cells [39, 40] were maintained at 5% CO_2_ and 37 °C in NFS-60-medium (RPMI 1640 medium with additional 1 mM L-glutamine, 1 mM Na-pyruvate, 10% FCS, 1% Antibiotic-Antimycotic (15240062, Gibco™) and 12.5% KMG-2: 5637-CM (Conditioned Medium from CLS Cell Lines Service)). Prior to the proliferation assay, the cells were washed 3 times with NFS-60-medium without KMG-2. The assay was performed in black 96-well plates (6005660, Perkin Elmer). For this purpose, 45.000 cells per well in a total volume of 150 µL NFS-60-medium without KMG-2 were cultured under maintenance conditions with different concentrations of designs or rhG-CSF for 48 h. Then, 30 µL of CellTiter-Blue® Reagent (G808, Promega) was added to each well and cells were further cultivated for approximately 90 min under maintenance conditions. Subsequently, the fluorescence of each well was recorded with a plate reader (Synergy H4 Hybrid Microplate Reader, Agilent BioTek) with an excitation wavelength of 560 nm and emission at 590 nm. The half-maximal effective concentration (*EC_50_*) of each design or rhG-CSF was determined by fitting the obtained fluorescence values to their corresponding concentrations using a four-parameter sigmoidal function and the Nelder-Mead Simplex algorithm from the Python module SciPy [41]. For the analysis of the maximum response of proliferative activity (*E_max_*), the mean of the three highest concentrations was estimated for each sample, such that all values considered for the analysis were in the activity plateau. These means were normalized to the maximum response within each experiment to account for the general variability of cell activity between the independent replicates. For statistical analysis, an ordinary one-way ANOVA followed by a Tukey HSD test was performed.

### Testing of granulopoietic activity of designed proteins in vivo

To test the effect of the designs on zebrafish granulopoiesis, equal volumes (4 nL) of Moevan_control (4 mg/mL), rhG-CSF (2 mg/mL), Boskar4 variants, bv6_st2 (2 mg/mL) and bv8_st2 (2 mg/mL) were injected into the cardinal vein of transgenic *Tg(mpx:GFP)* [42] larvae at 1.5-day post-fertilization (dpf). Injected larvae were incubated at 28 °C. To quantify neutrophils 24 hours upon injection, larvae were positioned and orientated laterally within cavities formed in 1% agarose on a 96-well plate and then imaged using an SMZ18 Nikon fluorescence stereomicroscope. The number of GFP-expressing neutrophils was automatically determined by Imaris software using the spot detection tool. Fold change of the neutrophils was calculated by normalizing to the average of uninjected mpx:*gfp* counterparts at the same developmental stage. Zebrafish lines were maintained according to standard protocols and handled in accordance with European Union animal protection directive 2010/63/EU and approved by the local government (Tierschutzgesetz §11, Abs. 1, Nr. 1, husbandry permit 35/9185.46/Uni TÜ). All experiments described in the present study were conducted on larvae younger than 5 dpf.

For the mouse treatment experiments, B6.SJL-PtprcaPepcb/BoyCrl (Ly5.1) mice, aged between 6 to 8 weeks, were treated with intraperitoneal injections (i.p.) of rhG-CSF, or Boskar4 variants, bv6_st2 and bv8_st2. A concentration of 300 µg/kg was used for each protein injecting mice every second day for a total of five injections. Bone marrow cells of treated mice were isolated by flushing with a 22G syringe and subsequent filtering through a 45 µm cell strainer. After that, cells were counted and used for flow cytometry analyses on a FACS Canto II (BD) and using FlowJo (BD). Cells were stained with DAPI (#D9542, Sigma), anti-mouse CD45.1 PerCP (#110725, Biolegend), anti-mouse Ly6C AF488 (#128022, Biolegend), and anti-mouse Ly6G APC (#127614, Biolegend) antibody. Gates were set according to fluorescence minus one (FMO) control. Mice were maintained under pathogen-free conditions in the research animal facility of the University of Tübingen, according to German federal and state regulations (Regierungspräsidium Tübingen, M 05-20 G).

### Single molecule imaging and analysis

For cell surface labelling, G-CSFR was N-terminally fused to the ALFA-tag. Additionally, to assess the co-expression of JAK2ΔTK it was C-terminally fused to mEGFP. Both proteins were encoded on pSems vectors including the signal sequence of Igκ (pSems-leader) [43]. HeLa cells (ACC 57, DSMZ Germany) were cultured as previously described [44]. For transient transfection, cells were incubated for 4-6 h with a mixture of 150 mM NaCl, 10 μL of 1 mg/mL polyethylenimine (PEI MAX®, Polysciences 24765) and 1500 ng (ALFAtag-G-CSFR) and 2500 ng (JAK2ΔTK-mEGFP) of the desired constructs. Labeling, washing and subsequent imaging were performed after mounting the coverslips into custom-made incubation chambers with a volume of 1 ml. Cells were equilibrated in medium with FBS but lacking phenol red supplemented with an oxygen scavenger and a redox-active photoprotectant (0.5 mg/mL glucose oxidase (Sigma-Aldrich), 0.04 mg/mL catalase (Roche), 5% w/v glucose, 1 μM ascorbic acid and 1 μM methylviologene) to minimize photobleaching [45].

Selective cell surface receptor labeling was achieved by using anti-ALFAtag NBs, which were site-specifically labeled by maleimide chemistry via a single cysteine residue at their C-termini [45]. NBs labeled with Cy3B (DOL: 1.0) and ATTO 643 (DOL: 1.0) were added at concentrations of 3 nM each, at least 10 min before imaging. Coverslips were precoated with poly-l-lysine-graft-poly(ethylene glycol) to minimize unspecific binding of NBs and functionalized with RGD peptide for efficient cell adhesion [46].

Single-molecule imaging was carried out by dual-color total internal reflection fluorescence microscopy using an inverted microscope (IX71, Olympus) equipped with a spectral image splitter (DualView, Optical Insight) and a back-illuminated electron multiplied CCD camera (iXon DU897D, Andor Technology). Fluorophores were excited by simultaneous illumination with a 561-nm laser (CrystaLaser; approximately 32LWLcm^−2^) and a 642-nm laser (Omicron: approximately 22LWLcm^−2^). Image stacks of 150 frames were recorded for each cell at a time resolution of 32 ms per frame, with at least 10 cells recorded in each experiment. Ligands were incubated for 10 min before imaging. All imaging experiments were carried out at room temperature. Prior to image acquisition, the presence of JAK2ΔTK-mEGFP was confirmed through emission with a 488-nm laser (Sapphire LP, Coherent)

Dual-color time-lapse images were evaluated using an in-house developed MATLAB software (SLIMfast4C, https://zenodo.org/record/5712332) as previously described in detail [45]. After channel registration based on calibration with fiducial markers, molecules were localized using the multi-target tracking algorithm [47]. Immobile emitters were filtered out by spatiotemporal cluster analysis [48]. For co-tracking, frame-by-frame co-localization within a cut-off radius of 100 nm was applied followed by tracking of co-localized emitters using the utrack algorithm [49]. Molecules co-diffusing for 10 frames or more were identified as co-localized. Relative levels of co-localization were determined based on the fraction of co-localized particles [46]. Diffusion properties were determined from pooled single trajectories using mean squared displacement analysis for all trajectories with a lifetime greater than 10 frames. Diffusion constants were determined from the mean squared displacement by linear regression.

### Analysis of STAT3/5 phosphorylation by Western Blot

NFS-60 cells were washed two times with NFS-60-medium (RPMI 1640 medium with additional 1 mM L-glutamine, 1 mM Na-pyruvate, 10% FCS, 1% Antibiotic-Antimycotic (15240062, Gibco™)) and starved at 300,000 cells/mL for 16 h at 5 % CO_2_ and 37 °C. Subsequently, the starved cells were treated with 1 nM of rhG-CSF or the selected design for 5 min or 30 min.

Following the treatment, the cells were consistently maintained at low temperatures on ice. They were harvested using a refrigerated centrifuge operating at 4 °C at 300 g for 2 min. The cells were washed once with ice-cold PBS and the dry pellets were frozen at – 70 °C until further use. Whole-cell lysates were obtained by lysing 1×10^6^ cells in 200 μL Laemmli buffer (30% glycerol, 6% SDS, 7.5% β-Mercaptoethanol, 0.75% Bromphenol blue in 200 nM Tris-HCL [pH 6.8]), which were subsequently heated at 95 °C for 5 min and spun down. Proteins were separated by dodecyl-sulfate polyacrylamide gel electrophoresis and transferred to Amersham™ Protran® nitrocellulose membranes or PVDF membranes (Invitrogen). The membranes were blocked with 5% non-fat dry milk-TBST (10 mM Tris-HCL [pH 8.0], 150 mM NaCl, 0.1% Tween® 20) for 1 h at room temperature and subsequently incubated with primary antibodies in 8 mL milk-TBST buffer overnight at 4 °C or for 2 h at room temperature (RT). After washing 4 times for 5 min with TBST, membranes were incubated with secondary antibodies in 8 mL milk-TBST buffer for 1 h at RT and then again washed 4 times for 5 min with TBST. The protein bands were detected using Pierce (Thermo Fisher Scientific) or Clarity (BioRad) ECL western blotting substrate kits, and imaged using Fusion FX (Vilber) or Sapphire™ (Azure Biosystems) imagers. The following antibodies and corresponding dilutions were used: primary rabbit polyclonal antibodies against: STAT5 1:500 (Cell Signaling, #9363S), phosphorylated STAT5 (Tyr694) 1:750 (Cell Signaling, #9351S), STAT3 1:200 (SantaCruz, # SC-482), alpha-tubulin 1:1000 (Cell Signaling, #2144S), primary rabbit monoclonal antibodies against: STAT5 1:1000 (Cell Signaling, #94205S), STAT3 1:1000 (Cell Signaling, #12640S), phosphorylated STAT3 (Tyr705) 1:1000 (Cell Signaling, #9145L), and GAPDH 1:1000 (Cell Signaling #2118S), and secondary horseradish peroxidase-conjugated anti-rabbit antibodies 1:3000 (Cell Signaling, #7074, or Jackson ImmunoResearch, #111-035-003). Gel images were exported for analysis as .tiff files and processed using FIJI software [50]. Following brightness/contrast adjustments, the bands of interest were isolated with the rectangle tool, and the peaks of band intensity were plotted. The peaks were individually separated using the line tool, and the areas under each peak were measured to determine the signal intensities of the immuno-detected proteins. Subsequently, these signal intensities were normalized by the corresponding signal of the housekeeping protein.

### Evaluation of time-dependent effects of the designed agonists on the proliferation of CD34^+^ HSPCs

Cells were incubated in poly-L-lysine–coated 96-well plates (2 × 104 cells/well) in Stemline II Hematopoietic Stem Cell Expansion medium (Sigma Aldrich; #50192) supplemented with 10% FBS, 1% penicillin/streptomycin, 1% L-glutamine and 50 ng/mL SCF, 20 ng/mL IL-3 and 10 ng/mL rhG-CSF, or designed agonists at different concentrations. using an IncuCyte S3 Live-Cell Analysis System (Essen Bio) with a 10 x objective at 37 °C and 5% CO_2_. Cell proliferation over time was analyzed using IncuCyte S3 Software. Experiments in this study involving human samples were conducted according to Helsinki’s declaration, and study approval was obtained from the Ethical Review Board of the Medical Faculty, University of Tübingen.

### Colony-forming unit (CFU) assay with human HSPCs

Human cord blood CD34^+^ cells were isolated from the mononuclear cell fraction by Ficoll density gradient centrifugation with subsequent magnetic bead separation using the Human CD34 Progenitor Cell Isolation Kit (Miltenyi Biotech Germany; #130-046-703). 1 × 10^4^ cells/mL CD34^+^ cells were plated in 35 mm cell culture dishes in 1 mL Methocult H4230 medium (Stemcell Technologies) supplemented with 2% FBS, 10 µg/mL 100x Antibiotic-Antimycotic Solution (Sigma), 50 ng/mL SCF, 20 ng/mL IL-3 and 20 ng/mL rhG-CSF, or 100 ng/mL designed agonists. Cells were cultured at 37 °C and 5% CO_2_. Colonies were counted on day 14.

### RNA-sequencing analysis

NFS-60 cells were washed two times with NFS-60-medium (RPMI 1640 medium with additional 1 mM L-glutamine, 1 mM Na-pyruvate, 10% FCS, 1% Antibiotic-Antimycotic (15240062, Gibco™)) and starved at 300,000 cells/mL for 16 h at 5% CO_2_ and 37 °C. 1.5 million cells per sample in a T25 cell-culture flask were stimulated with 1 nM of the selected design or rhG-CSF for 8 h. Subsequently, the cells were washed twice with PBS, lysed in 350 µL RTL buffer and stored at -70 °C until RNA extraction. At least 500 ng of RNA per sample was used to prepare libraries for RNA sequencing. The RNA Integrity Number (RIN) for RNA quality assessment was measured using a 2100 Bioanalyzer (Agilent). All RNA samples showed RIN > 9,6 (max = 10), demonstrating high quality of the RNA samples. Directional mRNA library preparation (poly A enrichment) and sequencing were done by Novogene (https://www.novogene.com). The libraries were sequenced on an Illumina NovaSeq 6000 using paired-end 150 bp sequencing mode with a minimum of 6 Gb of raw data per sample. For data analysis, nf-core/rnaseq (https://nf-co.re/rnaseq, [51]) was used, an RNA sequencing analysis pipeline that takes a sample sheet and FASTQ files as input, performs quality assessment, trimming, alignment, and produces a gene count (Table S1) and extensive QC report. Nf-core/rnaseq pipeline was run with the following parameters: --profile ‘docker,’ --genome ‘GRCm38,’ --aligner ‘star_salmon,’ pipeline version 3.10. Raw sequencing read counts were analyzed with the R package edgeR v3.42.2 [52]. 11,801 of initial 45,706 transcripts showed sufficient level of expression. Normalized log2 cpm values were used for PCA and descriptive part of the analysis. Differential expression was determined by fitting a quasi-likelihood negative binomial generalized log-linear model to count data. Resulting candidate genes that passed thresholds of absolute log2 fold change > 1 and FDR < 0.05 were tested for gene set enrichment using the R package clusterProfiler v4.8.1 [53]. Two gene set collections were used: GO terms and MSigDB m2 mouse collection v2022.1.

## Results

### Computationally-guided screening identifies higher-affinity G-CSFR binders

G-CSF dimerizes G-CSFR subunits in a 2:2 stoichiometry through two (high- and low-affinity) binding sites (Fig. 1A). Previously, we *de novo* designed a G-CSFR-binding module (Boskar4), that when tandemly linked (i.e. Boskar4 short tandem of <u>2</u> domains; B4_st2) could dimerize and activate G-CSFR (Fig. 1B) [33]. In contrast to G-CSF, Boskar4 binds G-CSFR through only one binding site that engages the CRH domain (Fig. 1C). Thus, our first aim was to enhance the affinity of the Boskar4 binding module to G-CSFR, which we reasoned could lead to more potent dimerization by the respective design agonists (i.e. B4_st2 variants). We built a computationally-guided mutant library by modelling the Boskar4:G-CSFR complex (Fig. 1C, D) and scanning for energy-minimizing Boskar4 mutations at the binding interface, using Damietta software [37]. In addition to the computationally-designed sequence library, we also built a site-saturation mutagenesis of the same positions to serve as control (Fig. S1). The designed library contained 6 mutations per position, in contrast to an average of 12 mutations per position for the control library. We used a bacterial display screening system [35] in which the ligand is presented as an extracellular Intimin fusion on the surface of the bacterial particles. Upon adding fluorescently-labelled G-CSFR, we sorted the bacterial particles for receptor binding using fluorescence-activated sorting (FACS) (Supplementary Methods). We ran both libraries (saturation and designed) through five cycles of FACS enrichment under the same conditions, where shifts in the fluorescence of both populations were observed across the enrichment cycles (Fig. 2A). Forty-eight single clones were picked from either library after the third and fifth FACS cycles, amounting to a total of 192 clones. Using on-plate fluorescence- and luminescence-based assays we quantified the relative binding of these clones to the receptor under the same conditions (Fig. S2).

**Figure 1.**
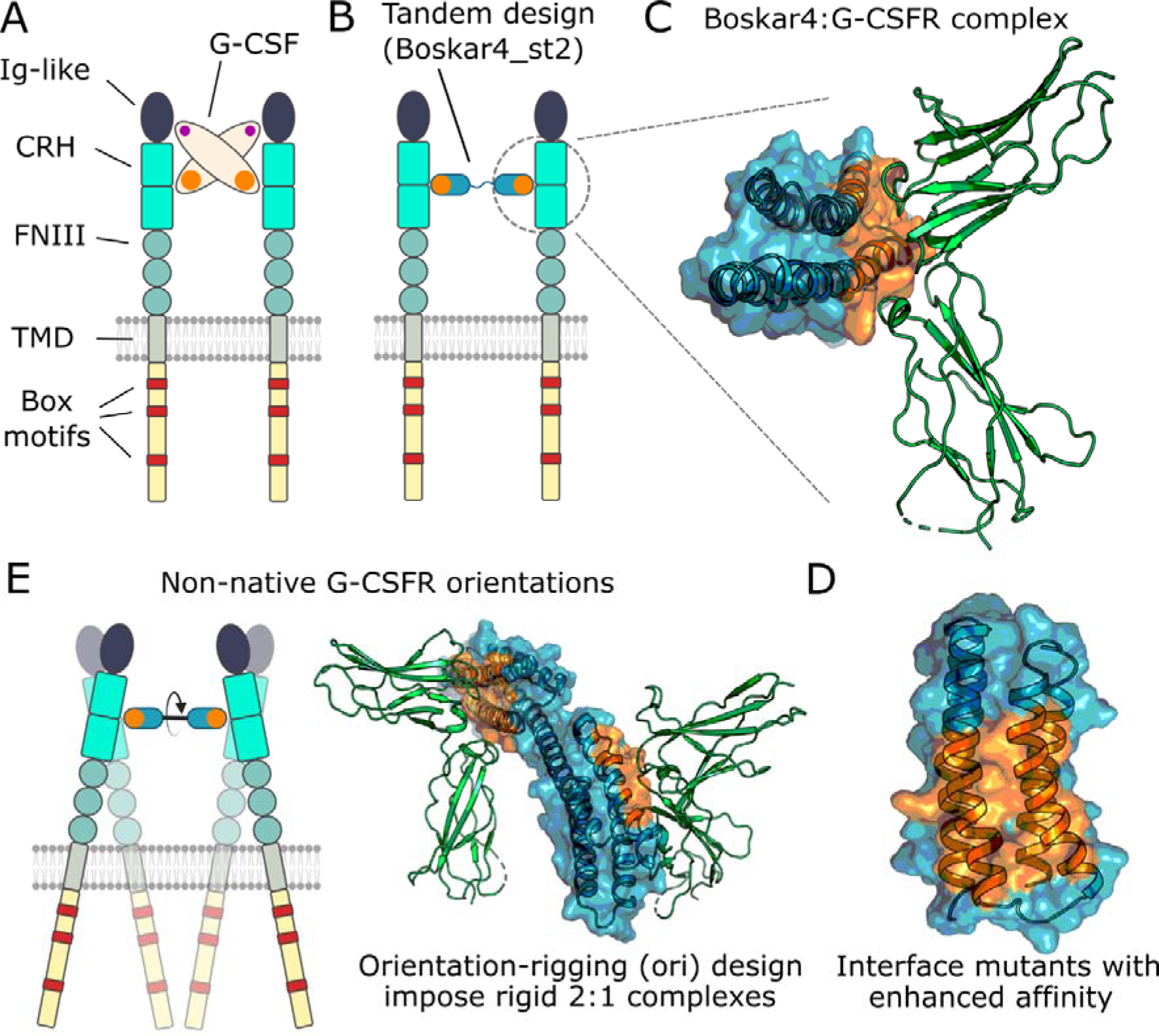
Overall strategy to generate novel G-CSFR modulators with varying affinities and geometries. **(A)** G-CSF (beige oval) activates G-CSFR by dimerizing the Ig-like and CRH domains (dark blue and cyan) through two distinct receptor-binding sites (orange and purple) [80]. **(B)** In contrast, the *de novo*-designed Boskar4_st2 agonist consists of two tandemly repeated Boskar4 domains, which **(C)** possess minimal architecture and encode a single high-affinity, receptor-binding site (blue domain and orange surface patch). **(D)** On the Boskar4 module, the residues forming the receptor-binding entity were diversified in order to identify affinity-enhanced variants. **(E)** The improved Boskar4 module was fused into rigid, orientation-rigging designs (oris) that can dimerize and activate G-CSFR in non-native geometries.

**Figure 2.**
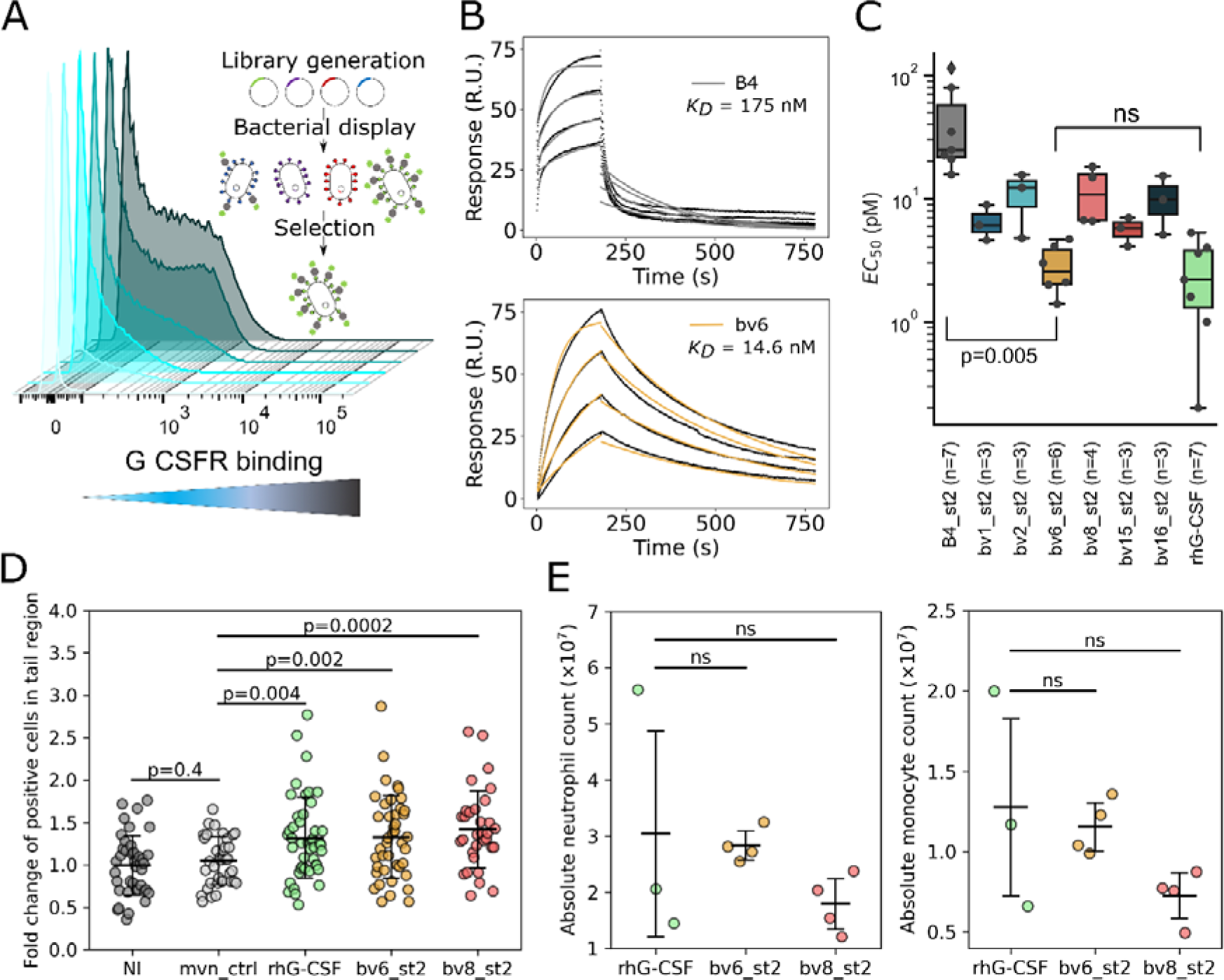
Affinity-enhanced binders exhibit improved granulopoietic activity *in vitro* and *in vivo*. **(A)** Cell sorting of bacterially-displayed Boskar4 mutants bound to fluorescently labeled G-CSFR was conducted over 5 steps of enrichment. Successive fluorescence shifts are shown as light cyan to dark gray distributions. **(B)** SPR titrations show single-domain bv6 to bind 12-fold tighter than its Boskar4 counterpart (Fig. S7D and Table 1). **(C)** Proliferative activity assays in NFS-60 cells showed the enhanced activity (*EC_50_*) of all variants in comparison to Boskar4_st2, where the most active variant bv6_st2 was not significantly different in activity compared to the native ligand (rhG-CSF; lightgreen box). For statistical analysis a one-way ANOVA over all samples was performed followed by a Tukey HSD test. Proliferation assays were conducted in at least three (3 < n < 7) independent experiments per variant or rhG-CSF (Table 1 and Fig. S8). **(D)** Interval plots of the fluorescent neutrophils in transgenic *Tg:mpxGFP* zebrafish larvae that were either not injected (NI), injected with the inactive protein Moevan_control (mvn_ctrl) [54], rhG-CSF, or indicated Boskar4 variants (bv6_st2 or bv8_st2) after 24 h of treatment. Data shows mean ± standard deviation, where each circle represents one zebrafish larva (NI n=40, mvn_ctr n=35, rhG-CSF n=44, bv6_st2 n=42, bv8_st2 n=32). Statistical analysis was carried out by ordinary one-way ANOVA followed by a Tukey HSD test. **(E)** C57BL/6 Ly5.1 mice were treated (i.p.) with rhG-CSF, bv6_st2, or bv8_st2 (each circle indicates one mouse). Absolute cell counts of mouse neutrophils (DAPI^-^CD45^+^Ly6G^+^Ly6C^+^, left image) and monocytes (DAPI^-^CD45^+^CD19^-^CD3^-^CD11b^+^Ly6C^+^Ly6G^-^, right image) in the bone marrow of treated mice are shown. Statistical analysis was carried out by ordinary one-way ANOVA with a multiple comparisons test of all experimental drugs to the G-CSF as a positive control (rhG-CSF n=3, bv6_st2 n=4, bv8_st2 n=4; ns: not significant).

**Table 1.** Binding affinity and activity parameters determined for G-CSFR modulators. The association rate constant (*k_a_*), the dissociation rate constant (*k_d_*) and the apparent equilibrium dissociation constant (*K_D_*) were obtained by surface plasmon resonance (SPR) against immobilized hG-CSF receptor (compare Fig. S6, S7, and S14). The mean and standard deviation (sd) of the half-maximal effective concentration *EC_50_* (pM) was obtained from NFS-60 activity assays of at least 3 biological replicates (compare Fig. S8 and S15).

| | $k_a$ ($M^{-1} s^{-1}$) | $k_d$ ($s^{-1}$) | Apparent $K_D$ (M) | Chi<sup>2</sup> (RU<sup>2</sup>) | EC<sub>50</sub> (pM) mean ± sd |
| --- | --- | --- | --- | --- | --- |
| **Boskar4** | $(3.4 \pm 1.7) \times 10^4$ | $(5.4 \pm 1.4) \times 10^{-3}$ | $(1.7 \pm 0.4) \times 10^{-7}$ | 13.9 | NA |
| **bv6** | $(2.4 \pm 0.7) \times 10^5$ | $(3.3 \pm 0.4) \times 10^{-3}$ | $(1.4 \pm 0.2) \times 10^{-8}$ | 2.51 | NA |
| **Boskar4_st2** | $(2.3 \pm 1.0) \times 10^6$ | $(3.3 \pm 0.7) \times 10^{-3}$ | $(1.6 \pm 0.3) \times 10^{-9}$ | 0.678 | 33.1 ± 23.6 |
| **bv1_st2** | $(9.7 \pm 2.0) \times 10^5$ | $(2.2 \pm 0.2) \times 10^{-4}$ | $(2.4 \pm 0.6) \times 10^{-10}$ | 1.44 | 6.5 ± 2.2 |
| **bv2_st2** | $(3.5 \pm 1.3) \times 10^5$ | $(1.7 \pm 0.1) \times 10^{-4}$ | $(5.3 \pm 1.8) \times 10^{-10}$ | 0.371 | 10.9 ± 5.5 |
| **bv6_st2** | $(4.9 \pm 0.9) \times 10^5$ | $(5.9 \pm 0.5) \times 10^{-4}$ | $(6.4 \pm 1.0) \times 10^{-10}$ | 1.16 | 2.9 ± 1.3 |
| **bv8_st2** | $(1.8 \pm 0.4) \times 10^6$ | $(1.5 \pm 0.1) \times 10^{-3}$ | $(8.3 \pm 1.6) \times 10^{-10}$ | 1.36 | 11.6 ± 5.9 |
| **bv15_st2** | $(7.4 \pm 5.0) \times 10^5$ | $(4.0 \pm 0.1) \times 10^{-4}$ | $(6.5 \pm 2.1) \times 10^{-10}$ | 0.982 | 5.6 ± 1.5 |
| **bv16_st2** | $(8.1 \pm 0.1) \times 10^5$ | $(7.1 \pm 0.8) \times 10^{-4}$ | $(8.8 \pm 1.2) \times 10^{-10}$ | 0.473 | 10.1 ± 5.1 |
| **rhG-CSF (self-made from *E. coli*)\*** | $(3.0 \pm 0.3) \times 10^5$ | $(4.9 \pm 2.8) \times 10^{-4}$ | $(1.1 \pm 1.6) \times 10^{-9}$ | 4.9 | 2.6 ± 1.8 |
| **Lenograstim (Glycosylated, commercial rhG-CSF)** | NA | NA | NA | NA | 1.0 ± 0.7 |
| **ori0** | $(6.5 \pm 6.6) \times 10^5$ | $(2.5 \pm 0.8) \times 10^{-4}$ | $(5.6 \pm 2.4) \times 10^{-10}$ | 1.11 | 2.8 ± 1.4 |
| **ori1** | $(1.4 \pm 2.1) \times 10^6$ | $(1.2 \pm 0.3) \times 10^{-4}$ | $(3.6 \pm 2.8) \times 10^{-10}$ | 0.779 | 28.9 ± 0.7 |
| **ori2** | $(1.1 \pm 0.8) \times 10^5$ | $(4.6 \pm 1.3) \times 10^{-5}$ | $(3.4 \pm 1.6) \times 10^{-10}$ | 2.83 | 35.6 ± 19.8 |
| **ori3** | $(8.7 \pm 5.9) \times 10^5$ | $(2.5 \pm 0.1) \times 10^{-4}$ | $(4.1 \pm 0.1) \times 10^{-10}$ | 2.36 | 10.6 ± 3.0 |
| **ori4** | $(1.2 \pm 0.1) \times 10^5$ | $(7.6 \pm 1.4) \times 10^{-5}$ | $(6.4 \pm 1.14) \times 10^{-10}$ | 1.37 | 19.3 ± 6.5 |

\* The SPR parameters of rhG-CSF (self-made purified from *E. coli*) were taken from Skokowa *et al.* 2022 [33].

The sequences of the top 40 clones picked from both libraries after Sanger sequencing yielded 16 unique Boskar4 variants (Fig. S2D). A focused FACS-based analysis for these ligands’ expression level and binding affinity on the surface of *Escherichia coli* (*E. coli)* (Fig. S3) showed the expression-normalized binding affinities to range from 3- to 37-fold increase, when compared to the starting Boskar4 clone. To narrow down the number of candidates for subsequent investigations, while keeping diversity, we chose six variants from the design library based on their estimated affinity, frequency in the screen, or the respective mutation composition (Fig. S2D and S3B). The most frequent variant was bv1, while the highest affinity variant was bv2. Four additional variants, bv6, bv8, bv15 and bv16 were chosen based on their middle- or lower-binding affinity profiles. Specifically, variants with polar and charged residues were included with the goal of selecting the most soluble and specific binders.

We constructed agonists by creating short-tandems of these six selected designs (Fig. S4), which were expressed and purified from *E. coli* and evaluated by size exclusion chromatography and SDS-PAGE analysis (Fig. S5). We used surface plasmon resonance (SPR) to determine the binding affinity and kinetic parameters of the designs to G-CSFR. The SPR results showed that all the six tested variants in their tandem forms bind G-CSFR within the sub-nanomolar affinity range, with up to 7-fold enhanced affinity compared to the starting template B4_st2 (Fig. S6 and Table 1). The observed affinity improvement was also higher when evaluated for single-domain variants. For instance, the single-domain form (bv6) of the most biologically active agonist (bv6_st2) exhibited a K_D_ of 14 ±2 nM, in comparison to the single-domain Boskar4 template, with K_D_ of 173 ±43 nM (Fig. 2B, S7B). Notably, circular dichroism (CD) measurements of bv6 showed the expected strong alpha-helical signal. Nanoscale differential scanning fluorimetry (nanoDSF) measurements demonstrated that bv6 exhibited hyper-thermostability, similar to Boskar4 (Fig. S7C, D).

### Affinity enhancement yields highly potent G-CSFR agonists with uncompromised thermostability

To evaluate the biological impact of increased affinity, we evaluated the proliferative activity of our design agonists in G-CSF-responsive NFS-60 cells [39, 40]. All the affinity-enhanced agonist variants induced stronger proliferation than the starting template (B4_st2). Remarkably, the most active design, bv6_st2 had an *EC_50_* of 2.9 ±1.3 pM compared to 33 ±23.6 pM for B4_st2 and 2.6 ±1.8 pM for rhG-CSF (Fig. 2C, S8 and Table 1), whereas no differences in the maximum response were noticed among the designs and rhG-CSF (Fig. S8E). Furthermore, the tested agonists showed no strong correlations between the proliferative activity (*EC_50_*) and the binding affinity (*K_D_*), the association rate constant (*k_a_*), or the dissociation rate constant (*k_d_*) (Fig. S9). We therefore expect that the biological activity within this narrow range of affinities might be influenced by other factors such as relative stabilities and binding specificity of the different agonists.

CD spectra of all design agonists indicated their strong alpha-helical nature (Fig. S10A), while nanoDSF melting curves highlighted their thermostability, where half of the tested variants (bv1_st2, bv6_st2 and bv8_st2) exhibited no detectable unfolding up to 110 °C, while the other three (bv2_st2, bv15_st2 and bv16_st2) exhibited a melting onset at around 100 °C. None of the design agonists exhibited increased scattering over the full temperature range (Fig. S10B-H), highlighting their excellent colloidal stability. Similar results were observed for the single-domain form of the designs (Fig. S7D).

These results indicate, that the affinity-driven functional enhancement did not alter the biophysical properties of the designs, which are far superior to rhG-CSF. Therefore, we further sought to test whether the design agonists exhibit similarly enhanced activity *in vivo*.

### Boskar4 variants induce neutrophil production in zebrafish embryos and mice

We first tested the capacity of two design agonists, bv6_st2 and bv8_st2 to induce granulopoiesis in a fluorescent neutrophil reporter zebrafish transgenic line, *Tg(mpx:GFP)* [42]. Interestingly, the increased number of GFP^+^ neutrophils localized near the caudal hematopoietic tissue (tail region) was comparable between larvae treated with rhG-CSF and bv6_st2 or bv8_st2 24 h post-injection (Fig. 2D). The number of neutrophils did not significantly increase in the Moevan_control-injected larvae. Moevan is an inert helical protein [54] expressed and purified under the same conditions.

We additionally treated C57BL/6 Ly5.1 mice with 300 μg/kg of rhG-CSF, bv6_st2, or bv8_st2 by intraperitoneal (i.p.) injection every second day with five injections in total. One day after the fifth injection, the number of Ly6G^+^Ly6C^+^ neutrophils and Ly6G^-^Ly6C^+^ monocytes in the bone marrow of treated mice was evaluated. The treatment of mice with bv6_st2 or bv8_st2 induced the production of neutrophils and monocytes at a comparable level to rhG-CSF (Fig. 2E). These results confirm the designs capacity to activate G-CSFR *in vivo*.

### Design of agonists altering the relative orientation of G-CSFR

Besides affinity, the geometry of receptor dimerization can also impact its activation. Hence, we sought to design orientation-rigging (ori) agonists that are composed of two rigidly-connected bv6 modules (Fig. 1E). We thus modelled helical connectors to introduce different inter-domain rotations (Fig. S11A, B), and predicted the 2:1 assemblies using AlphaFold2 [55] (Fig. 3A). The five templates named ori0-4 underwent sequence design using the combinatorial sampler application of the Damietta software (v0.36) [37], and filtering based on their structural rigidity in molecular dynamics simulations (Fig. S11C, D). Our models predicted spacings across the receptor transmembrane domains (TMD) of *d_0_* = 78 ±32 Å, *d_1_*= 206 ±14 Å, *d_2_* = 287 ±6 Å, *d_3_*= 155 ±35 Å, and *d_4_* = 143 ±37 Å, for ori0 through ori4, respectively, which compares to *d_g_* = 57 Å for the native G-CSF:G-CSFR complex (Fig. 3A, B).

**Figure 3.**
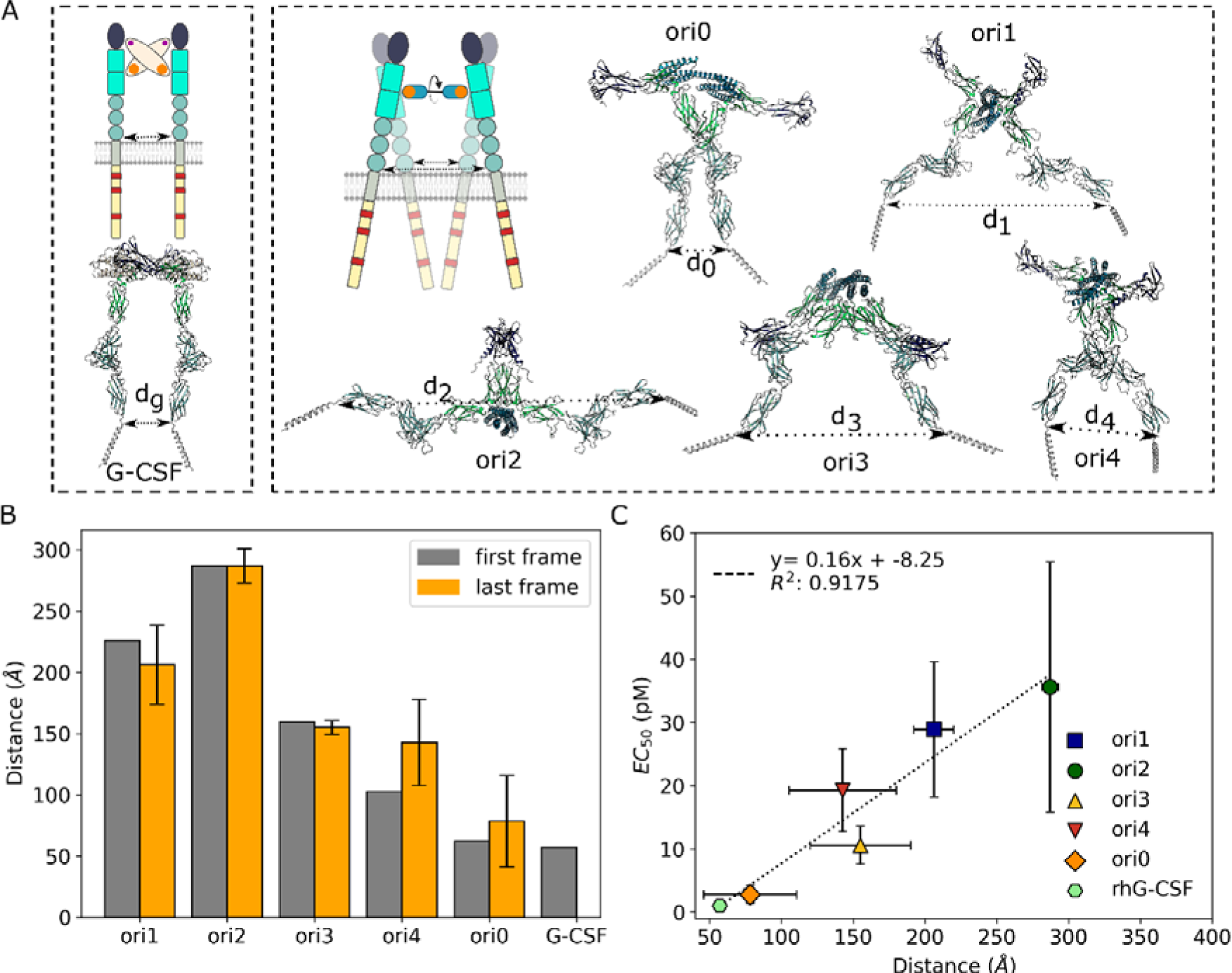
Agonists designed to associate G-CSFR in non-native dimeric geometries yield distinct cell activation potencies. **(A)** Unlike the native 2:2 ligand-receptor assembly induced by G-CSF (PDB: 2D9Q), we developed a range of novel ligands by rigidly connecting two copies of the affinity-enhanced bv6 module (Supplementary Methods; Fig. S11) that form 1:2 ligand-receptor complexes. Modeled complexes of these orientation-rigging designs (ori0 through ori4) and G-CSFR indicate varying transmembrane domain spacings (*d_x_*). Ori0 was designed to closely mimic this distance parameter to that of G-CSF (i.e., *d_0_* ≈ *d_g_*). **(B)** Molecular dynamics simulations performed on a selected set of designed ori candidates confirmed the rigidity of the design linkers. Orange bars represent the mean spacing and its standard deviation from 5 simulation replicas. **(C)** Proliferation assays in NFS-60 cells of the five selected ori designs and rhG-CSF (Lenograstim) showed the proliferative *EC_50_*(mean and standard deviation of at least 5 independent experiments) to strongly correlate to the modelled transmembrane domain spacing (*d_x_*) for the different proteins. The dotted line represents the linear fit between the obtained *EC_50_* and the distances of the last frame (where the *d_g_* of G-CSF was obtained as a single value from the crystal structure (PDB:2D9Q).

The produced ori designs were helical, stable, and monomeric, with the exception of ori1 which showed monomeric-dimer exchange (Fig. S12 and S13). SPR titrations showed the ori designs to bind G-CSFR as expected, with affinities ranging from 0.44 nM to 0.73 nM (Fig. S14 and Table 1).

### The geometric modulation of the G-CSFR by design agonists differentially affects cell proliferation

Proliferation assays in NFS-60 cells showed the ori designs *EC_50_*values to range from 36 ±19.8 pM down to 2.8 ±1.4 pM (Table 1 and Fig. S15), where only ori1 displayed a reduced maximum proliferation level (*E_max_*) compared to rhG-CSF (Fig. S15H). The proliferative *EC_50_* values of these design agonists correlated well with the modelled distances across the receptor TMD for these designs (*R^2^* = 0.92; Fig. 3C). Ori0 was the most potent as it dimerizes G-CSFR with similar TMD spacing to G-CSF (*d_0_* ≈ *d_g_*). Conversely, the design agonists inducing the farthest TMD spacing, ori1 and ori2, showed the lowest proliferation.

### Design agonists impose distinct complexes with G-CSFR and can override the native ligand

To query the properties of these design:receptor complexes in a cellular environment, we explored G-CSFR assembly in the plasma membrane by live cell single-molecule fluorescence imaging. To this end, we fused ALFA-tag to the N-terminus of G-CSFR to an, expressed it in HeLa cells, and labeled it by a mixture of anti-ALFA nanobodies conjugated with Cy3B and AT643 (Fig. 4A) [56]. We additionally transfected the cells with JAK2 C-terminally fused to mEGFP, as the latter was shown to contribute to receptors assembly [7]. This construct lacks the Tyrosine kinase domain (TK) to eliminate any bias from downstream signaling activation. Simultaneous dual-color single molecule imaging by total internal reflection fluorescence microscopy enabled identifying individual G-CSFR dimers by co-localization and co-tracking analysis (Supplementary Movie S1, Fig. 4B and S16A).

**Figure 4.**
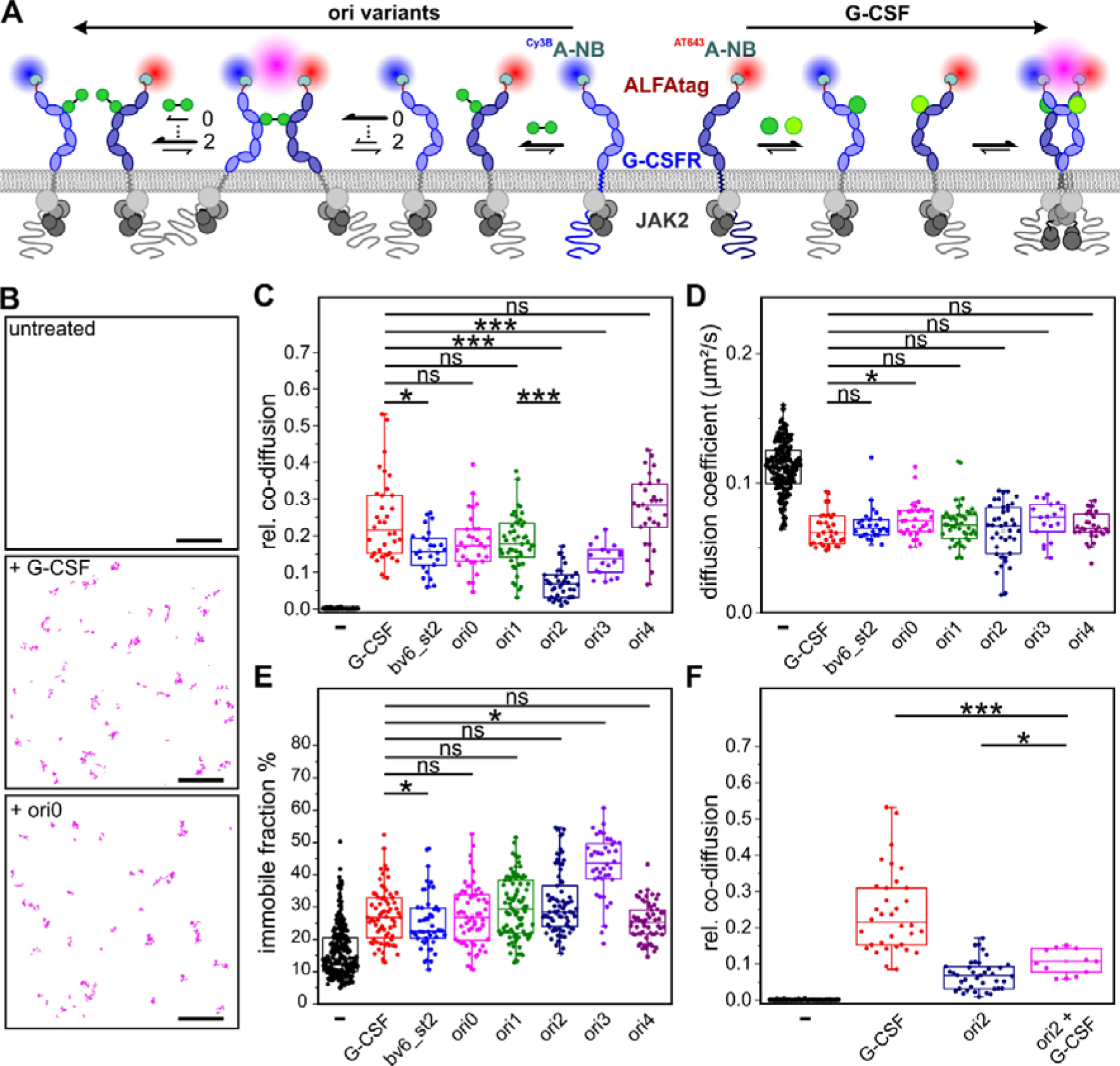
The design agonists differentially dimerize G-CSFR subunits in live cells. **(A, B)** Dual-color single molecule co-tracking for quantifying G-CSFR dimerization in live cells. **(A)** G-CSFR labeled via an N-terminal ALFA-tag using a mixture of nanobodies conjugated with Cy3B and AT643, respectively. The G-CSFR monomer-dimer equilibrium probed by dual-color single molecule co-tracking depends on the two-dimensional and three-dimensional binding affinities of the respective design agonist. Different dimerization principles by G-CSF (arrow to the right) and design agonists arrow to the left), as well as the consequences of altered orientation on the assembly kinetics are schematically outlined. (numbers on arrows indicate expected impact of ori0 (0) or ori2 (2) **(B)** Typical single molecule co-trajectories of G-CSFR in the absence of agonist and presence of G-CSF or ori0, respectively. A ROI from a single cell is shown. Scale bar: 5 µm. **(C)** Relative dimerization levels of G-CSFR in the absence of ligand and in the presence of G-CSF and design agonists. **(D)** Diffusion constants of G-CSFR monomers in the absence of ligand compared to G-CSFR dimers in the presence of G-CSF and design agonists identified by co-tracking analysis. **(E)** Comparison of the immobile fraction of G-CSFR in the absence of ligand and in the presence of G-CSF and design agonists. **(F)** Comparison of G-CSFR dimerization by G-CSFR by G-CSF, ori2 and an equimolar mixture of both. Box plots in C-F were obtained from analysis of multiple cells (For C-E; Numbers of cells and experiments, respectively, are 124 and 17, 34 and 3, 25 and 2, 30 and 2, 47 and 3, 39 and 3, 18 and 2, 30 and 2 for each column, going from left to right. For F; Numbers of cells and experiments, respectively, are 124 and 17, 34 and 3, 39 and 3, 10 and 1 for each column, going from left to right), with each dot representing the result from one cell. Box plots indicate data distribution of the second and third quartiles (box), median (line), mean (square) and 1.5× interquartile range (whiskers). Statistics were performed using two sided two-sample Kolmogorov-Smirnov test (not significant: ns;*P ≤0.05***P ≤ 0.001)

As expected, G-CSFR dimerization was negligible in the resting state, but strongly stimulated by G-CSF and all design agonists (Fig. 4C). While similar dimerization levels were observed for most design agonists, dimerization induced by ori2 and ori3 was significantly lower. Importantly, similar receptor cell surface densities were used in all experiments (Fig. S16B), thus ensuring that dimerization levels were not biased by the two-dimensional (2D) receptor concentration. Since the three-dimensional (3D) binding affinity of all design agonists is identical as they share the same bv6 binding site, the distinct differences in dimerization potencies of different design agonist can be ascribed by different 2D binding affinity (Fig. 4A). This can be rationalized by changes in the 2D association rate constant k^2D^, but not the complex stability k^2D^ due to geometrical constraints imposed by the orientation of the receptor ectodomains at the cell surface. With the formation of 1:1 G-CSFR:agonist complexes competing with G-CSFR dimerization, reduction in 2D binding affinity is accompanied by a lower maximum of the bell-shaped concentration-dimerization relationship of homomeric dimerizers [57]. Changes in dimerization efficiency can explain differences in the *EC_50_* values, as the overall receptor binding affinity depends on dimerization efficiency [58, 59], which was reported for synthetic ligands of EPOR [20, 21]. However, different G-CSFR dimerization levels induced by different design agonists did not directly correlate with their activity. For instance, ori1 showed substantially higher dimerization levels as compared to ori2, yet their NFS-60 proliferation activity was similar. Additionally, ori1 and ori0 had similar dimerization levels, but ori0 had about ten times higher proliferation activity compared to ori1 (Table 1). These results therefore confirm that biased signaling observed for our design agonists is related to the altered geometry of the signaling complex rather than weakened dimerization.

Our analysis of G-CSFR complexes indicated that their diffusion coefficients were similar among the different ligands, which were ∼45% lower than unbound G-CSFR monomers (Fig. 4D). However, the fraction of immobile G-CSFR increased by a factor of ∼2, with ori3 showing surprisingly higher immobilization by a factor of ∼3 (Fig. 4E). Receptor immobilization is very likely caused by endocytosis, and therefore differences across design agonist may indicate altered endocytic uptake [60]. Finally, we tested if the low-dimerizing ori2 can override native complex formation in the presence of G-CSF. We observed effective competition of G-CSF by ori2 at equimolar concentrations (Fig. 4F and S16C, D). Taken together, these results corroborate that our design agonists potently induce G-CSFR dimerization at the cell surface and that biased signaling can be explained by systematically altered complex geometries.

### Different design agonists tune G-CSFR signaling amplitudes, kinetics and cellular response patterns

Since STAT3 phosphorylation occurs at the membrane-distal regions of the intracellular segments of the receptor, while STAT5 is phosphorylated at the membrane-proximal receptor region [61], we reasoned that the relative ratio of pSTAT3/5 might be altered by different designs. Western blot analysis after stimulating NFS60 cells for 5 and 30 min confirmed increased phospho-STAT3 (pSTAT3, Tyr705) levels for bv6_st2 as compared to B4_st2, confirming the increased potency by enhancing the affinity (Fig. 5A, C and S17). For the oris however, striking differences were observed: while after 5 and 30 minutes ori0 and ori2 reached similar pSTAT3 levels and activation kinetics to bv6_st2, markedly lower pSTAT3 levels were found for ori1 for 5 min of stimulation (Fig. S17). Designed proteins showed substantially slower and prolonged STAT5 phosphorylation (pSTAT5, Tyr694) in comparison to rhG-CSF (Fig. 5B, D). Again, bv6_st2 and ori0 phosphorylated STAT5 more efficiently than B4_st2, ori2, or ori1, but the magnitude of pSTAT5 levels was markedly weaker for all variants compared to rhG-CSF. These results highlight that amplitudes and kinetics of STAT3 and STAT5 phosphorylation are affected by G-CSFR dimerization geometry.

**Figure 5.**
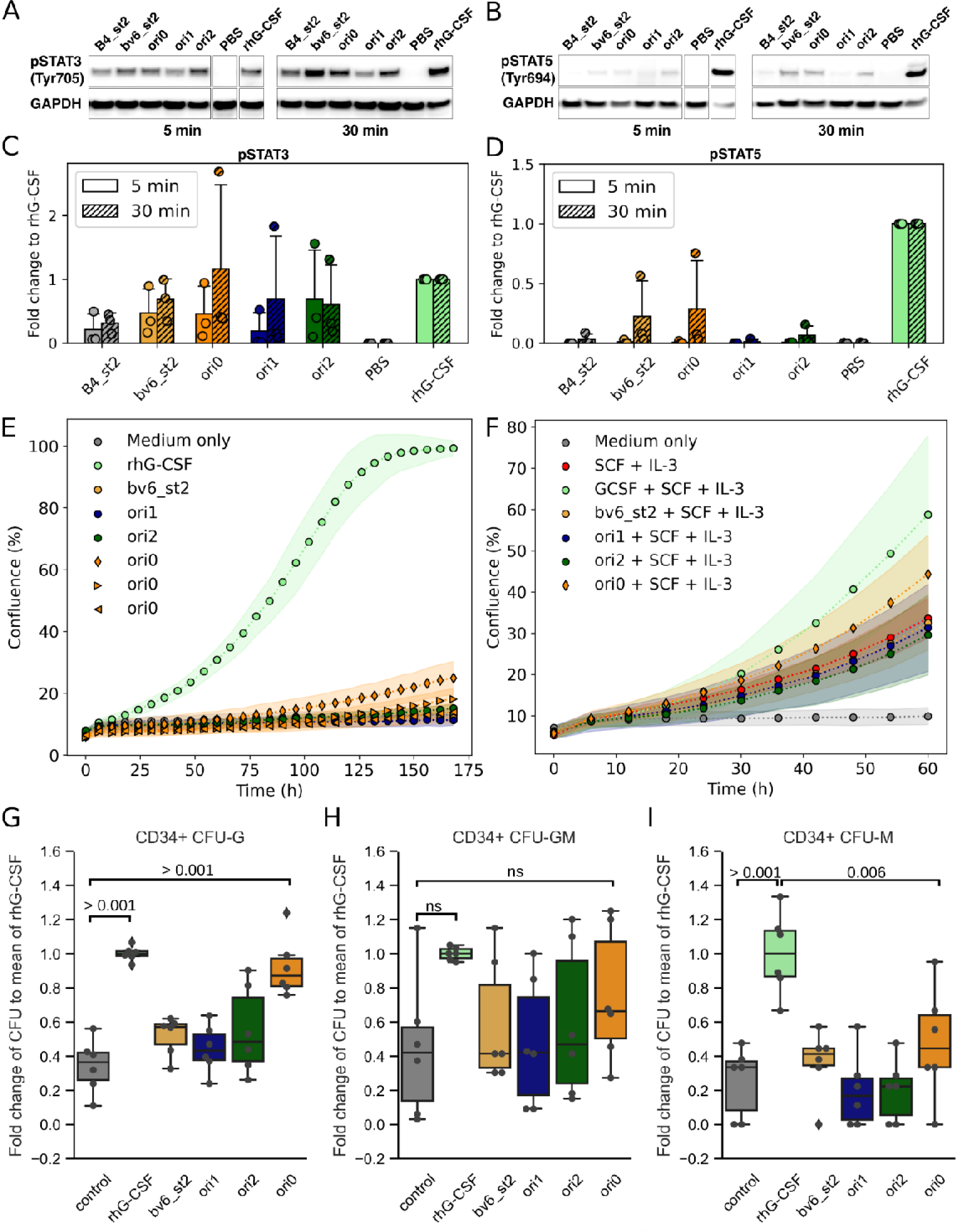
The affinity and geometry of the design agonists bias intracellular signaling and primary stem cell differentiation. **(A, B)** Representative western blot images of intracellular levels of phospho-STAT3 (Tyr705) and STAT3 (A) and phospho-STAT5 (Tyr694) and STAT5 (B) proteins after treatment of NFS-60 cells with 1 nM of different designs (saturating conditions) for 5 min (left pane) or 30 min (right pane). Glyceraldehyde 3-phosphate dehydrogenase (GAPDH) staining was used as a loading control. Three biological replicates of this experiment showed the same trend (Fig. S17). **(C, D)** The pSTAT3 and pSTAT5 levels were quantified after 5 min and 30 min from those replicates and are represented in (**C**) and (**D**), respectively. Shown is the fold change to the GAPDH normalized to the rhG-CSF treated samples. **(E, F)** To probe the effect of the agonist designs (100 ng/ml) and rhG-CSF (10 ng/ml) on healthy donoŕs CD34^+^ HSPCs, proliferation assays were performed without (**E**) or with the addition of 50 ng/mL SCF and 20 ng/mL IL-3 (**F**). Ori0 without SCF and IL-3 was tested at three concentrations 1 ng/mL (triangle-left), 10 ng/mL (triangle-right), and 100 ng/mL (diamond). Shown is the mean (points) and standard deviation (shades) of three parallel replicates. **(G, H, I)** Additionally, we performed CFU assays of healthy donoŕs CD34^+^ and progenits incubated on semi-solid medium supplemented with corresponding cytokines and design agonists. IL3 and SCF was added to all samples. Shown is the fold change to rhG-CSF of the quantified colony-forming units (CFU) of granulocytes (CFU-G) (**G**), granulocyte-macrophages (CFU-GM) (**H**), and macrophages (CFU-M) (**I**). The circles represent the obtained values for each condition of three independent experiments with two parallel replicates each. The indicated p-values were estimated by an ordinary one-way ANOVA followed by a Tukey HSD test.

G-CSF induces proliferation and granulocytic differentiation of HSPCs, and until now it is impossible to dissect these two cellular outcomes intentionally. Different magnitudes and kinetics of STAT3/STAT5 phosphorylation have different outcomes on HSPCs functions, where elevated pSTAT3 triggers myeloid differentiation, while elevated pSTAT5 induces cell proliferation [62]. We found that the designed proteins alone triggered only weak proliferation compared to rhG-CSF (Fig. 5E), which was potentiated by IL-3 and SCF (Fig. 5F). In line with previous observations, ori0 had the highest proliferative activity, but was still weaker than rhG-CSF. Nonetheless, ori0 was as active as rhG-CSF in inducing granulocytic differentiation of HSPCs when combined with IL-3 and SCF (Fig. 5G, H). Meanwhile, the other design agonists (ori1, ori2 and bv6_st2) did not significantly induce CFU-Gs in comparison with untreated cells. Interestingly, all the design agonists had a weak effect on the formation of monocytic colonies (CFU-M) (Fig. 5I), suggesting their selective activation of granulocytic differentiation, when compared to rhG-CSF. These results further establish the role of G-CSFR association geometry in fine-tuning of STAT3/STAT5 phosphorylation and dissecting proliferation from granulocytic differentiation of HSPCs.

### Designed G-CSFR agonists modulate intracellular transcriptional programs with hematopoietic bias

To assess the transcriptomic footprints of our design agonists, we carried out RNA-Seq of NFS-60 cells undergoing 8-h treatment with 1 nM of agonist or rhG-CSF. G-CSF regulated expression of 1950 genes compared to the PBS-treated control group (log FC 1, FDR less than 0.05). The number of differentially expressed genes was lower for all designs, and was affected by the inter-TMD spacing (e.g. 1292 genes for ori0 versus 524 genes for ori1) and binding affinity (e.g. 1115 genes for bv6_st2 versus 945 genes for B4_st2). Differentially regulated genes overlapped across all design agonists and were largely subsets of the rhG-CSF group (Fig. 6B, C). The general trend revealed a graded number of regulated genes across the groups, following the order G-CSF > ori0 > bv6_st2 > B4_st2 > ori2 > ori1 > PBS (Fig. 6C and Fig. S18). Gene set enrichment analysis (GSEA) showed similar enrichment patterns, particularly in the regulation of hematopoiesis/myelopoiesis-related gene sets among rhG-CSF and the design agonists, which followed the same trend (Fig. 6D and Fig. S19). As with individual genes, the degree of regulation of the enriched gene sets was either similar to the rhG-CSF group, or gradually reduced (Fig. S20).

**Figure 6.**
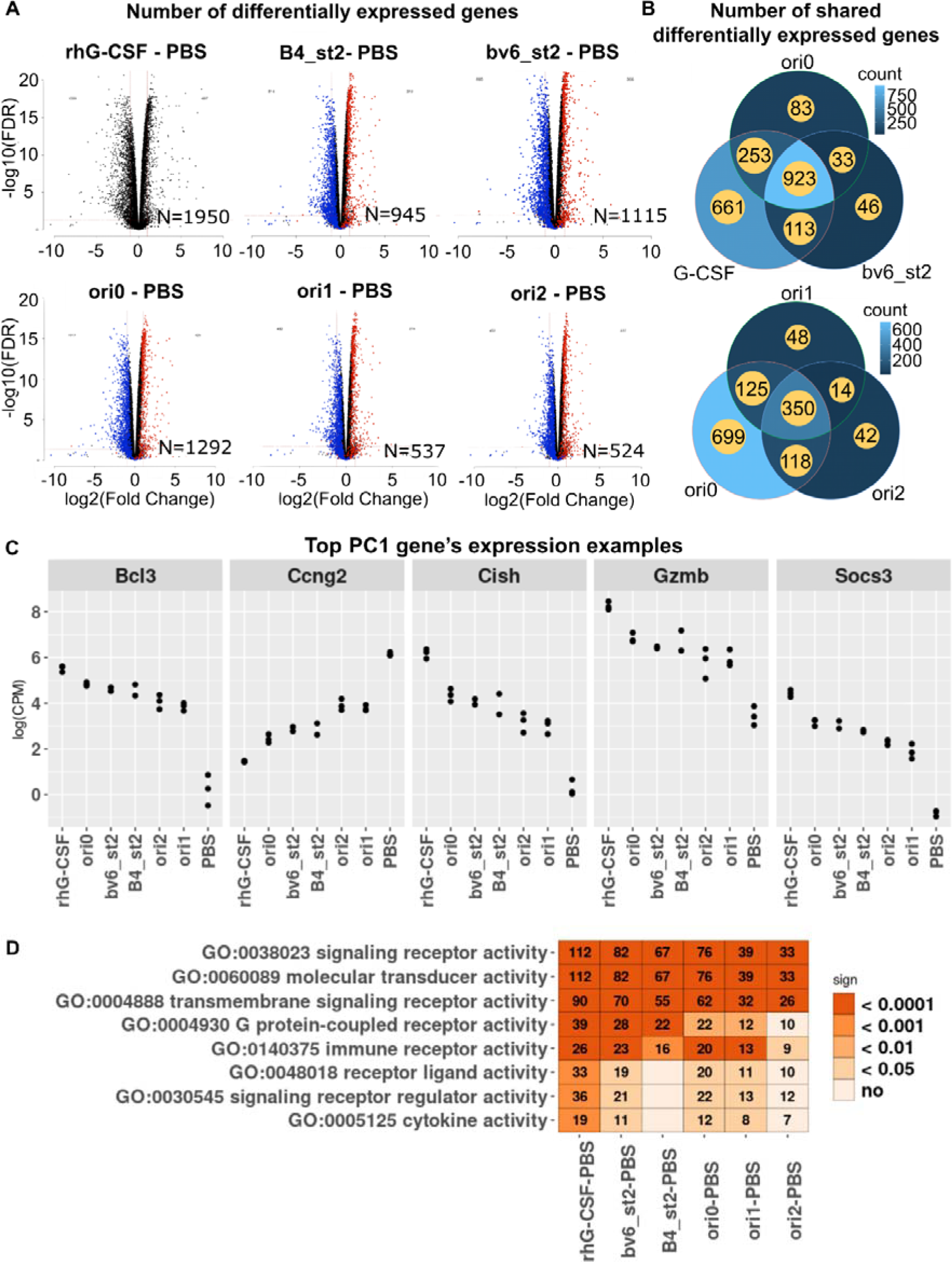
The design agonists regulate G-CSFR downstream signaling to varying degrees. **(A)** The number (N) of differentially expressed genes in NFS-60 cells after treatment with rhG-CSF or different designs compared to PBS treated control. Blue (downregulated genes) and red (upregulated genes) dots in the volcano plots indicate genes that are shared between the corresponding design and rhG-CSF. **(B)** The Venn diagrams show the numbers of shared and design-specific regulated genes. **(C)** Examples of the top five differentially expressed hematopoiesis-related genes out of 20 genes. **(D)** Gene Ontology (GO) analysis of significantly regulated signaling pathways in the indicated groups. Each number indicates the number of differentially expressed genes related to the corresponding pathway and the colours represent the significance score.

A large number of genes was exclusively regulated by rhG-CSF, including that involved in immunomodulation of T lymphocytes, myeloid-derived suppressor cells (MDSCs), monocytes/macrophages, dendritic cells and basophils, chemokine receptors (CXCR2, CXCR4, CX3CR1, CCR5) as well as VEGFR/ Kdr, transcription factors HOXA1, HOXB3, Klf3, Junb, MEIS3, Egr1, HOXC4, cell cycle regulators CDKN1A/P21 and CDKL2 and some adhesion molecules (Table S1).

## Discussion

D*e novo* design of cytokine receptor ligands with small and hyper-stable structures offers several advantages. This strategy gives room for functional enhancements that typically erode structural stability [63], and provides building blocks for constructing rigid larger structure to module receptors in novel ways. In this work, we leverage the *de novo*-designed G-CSFR binder (Boskar4) [33] through affinity and geometry engineering to tune G-CSFR activity by inducing specific signaling patterns and thus distinct cellular activity.

By enhancing the design’s receptor binding affinity, and thus dimerization efficacy, we could create agonists with increased granulopoietic activity *in vitro* and *in vivo*. The enhanced designs remained hyper-thermostable, in contrast to all previous studies aiming to improve the stability of rhG-CSF which only led to marginal improvements in its biophysical properties [64–66]. The design agonists stability and the diverse binding site sequences are invaluable for de-risking further optimization to cloak any immune epitopes during further pre-clinical development. After affinity, we sought to explore the role of receptor association geometry, whereby our results set forth the inter-TMD spacing G-CSFR subunits as another key parameter of signaling. The analysis of the phosphorylation of the main secondary messengers STAT3 and STAT5 revealed that design agonists with higher affinity and shorter inter-TMD spacing selectively increase the intracellular level of pSTAT3 while only minimally inducing pSTAT5. This might relate to the different localization of STAT3 and STAT5 along the intracellular segment of G-CSFR, and the mechanisms by which they are phosphorylated [61, 67, 68], where the inter-TMD spacing can affect the balance between these two processes. Broadly speaking, pSTAT3 primarily guides differentiation, while pSTAT5 mainly drives proliferation [67, 69–71]. Corresponding results were obtained through CFU assays in primary human HSPCs, where more affine and shorter-spacing designs led to more selective formation of granulocytic colonies, in comparison to G-CSF.

Single molecule fluorescence imaging confirmed the ability of most design agonists to potently dimerize G-CSFR at comparable levels to G-CSF. The reduced dimerization levels observed for ori2 and ori3 indicate that manipulating the geometry of G-CSFR dimers is accompanied by a loss of 2D association kinetics. Changes in signaling activity and specificity, however, cannot be attributed to changes in dimerization and rather support the idea that design agonists associate G-CSFR dimers with altered geometry. The very similar diffusion constants of all agonist-induced dimers, however, suggest more subtle changes in orientation as compared to the predicted structures, which should significantly reduce mobility. This could be explained by the additional strain imposed by membrane anchoring and additional interactions involving associated JAKs. Nonetheless, the distinct rise in the immobile fraction observed for ori3 points towards changes in endocytic uptake for that particular agonist. This warrants further investigation, as endocytic trafficking of cytokine receptor signaling complexes is emerging as a complex determinant regulating potency and kinetics of signal activation and the specificity of cellular responses [60, 72, 73]. Indeed, we observe distinctly reduced STAT5 phosphorylation magnitude and different kinetics of STAT3 phosphorylation for the different design agonists, which could be related to altered endocytic trafficking.

By looking at the gene expression footprints of our design:receptor constellations, we observed varying degrees of tuning of the G-CSFR-triggered transcription activation or inhibition, where the design agonists minimally affect the expression of genes that are not regulated by rhG-CSF. Thus, we could fine-tune G-CSFR signaling without regulating unexpected transcriptional pathways that are not linked to physiological G-CSFR activation. This information opens up an entirely new perspective in designing hematopoietic and granulopoietic proteins with specific activity and less unwanted functions, when compared to recombinant cytokines. To date, only *de novo* designed IL-2 and IL-4 mimics, that bind a subset of the natural cytokine’s heteromeric receptor subunits, could achieve more selective biological responses [74, 75]. Alternatively, tuning of homodimeric receptors, could rather be achieved through engineered agonists with varying affinities and geometries for EPOR [20, 21] and TPOR [76, 77]. Our results are first to establish the tunable modulation potential for G-CSFR.

These findings demonstrate the potential for tailoring G-CSFR agonists to achieve more specific therapies for hematopoietic stem cell disorders, and overcome the functional pleiotropy of G-CSF. In addition to inducing proliferation, migration, and myeloid differentiation of HSPCs, G-CSF also exerts anti-apoptotic and immunomodulatory activities [30, 61, 78, 79]. For instance, rhG-CSF’s widest use is to treat chemotherapy-induced neutropenia [31], whereby in some cases it may inadvertently activate G-CSFR-expressing tumor cells or dysregulate anti-cancer immune control of G-CSFR-expressing monocytes/macrophages, MDSCs, T-lymphocyte subsets, dendritic cells or natural killer cells. These adverse outcomes motivate the discovery of G-CSFR ligands capable of selective induction of specific granulopoietic signaling on HSPCs with minimal effect on proliferation or immune cell functions. This study thus paves the way for the further development of potent, hyperstable, and selective tunable sub-agonists of G-CSFR.

## Acknowledgements

This project has received funding from the IMPRS (TU, KM), the European Research Council under the European Union’s Horizon 2020 research and innovation program (grant agreement No 863952 (ACE-OF-SPACE)) (PM), the M. Schickedanz Kinderkrebsstiftung (ME, JS, NA), Deutsche Forschungsgemeinschaft (DFG; No 500215849; ME, BHA, JS, VH), (DFG; PI 405/15-2 – 326558201 and SFB; 1557, P13 – 467522186; JP), BMBF MyPred (JS) and the InnoChron COST EU action (JS). The authors would also like to thank Luis Ángel Fernández for providing the bacterial display system plasmid, and Regine Bernhard, Gabriele Hikade and Hella Kenneweg for technical assistance.

## Supplementary figures

**Supplementary Figure 3.**
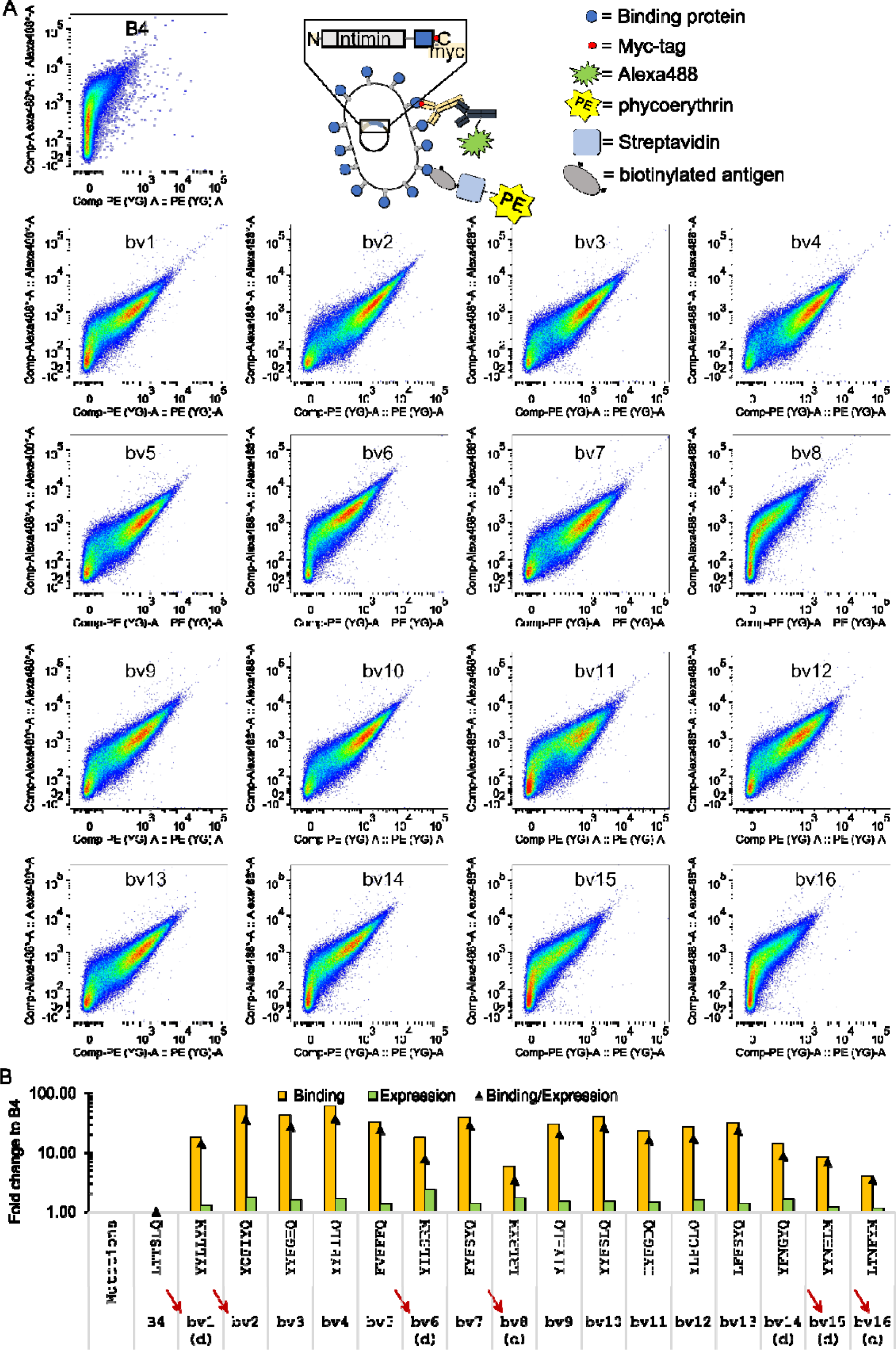
**(A)** The 16 new affinity-enhanced Boskar4 variants (bv1 to bv16) and Boskar4 (B4) were analyzed on the surface of *E. coli* by FACS for their binding activity to BioGCSFR (10 nM, PE channel) and the expression level of the corresponding variant (anti-myc labeling, Alexa488 channel). **(B)** Based on the measurements in (A) the binding to G-CSFR (orange bars), expression of the binder (green bars), and expression-normalized binding to G-CSFR (black triangles) of the affinity-enhanced Boskar4 variants normalized to Boskar4 was analyzed. Sequences selected for further characterization are marked with dark red arrows. The small “d” in parentheses (d) indicates that the variant was selected from the designed library. Additionally, the corresponding mutation set of each enhanced variant is indicated in comparison to Boskar4.

**Supplementary Figure 4.**
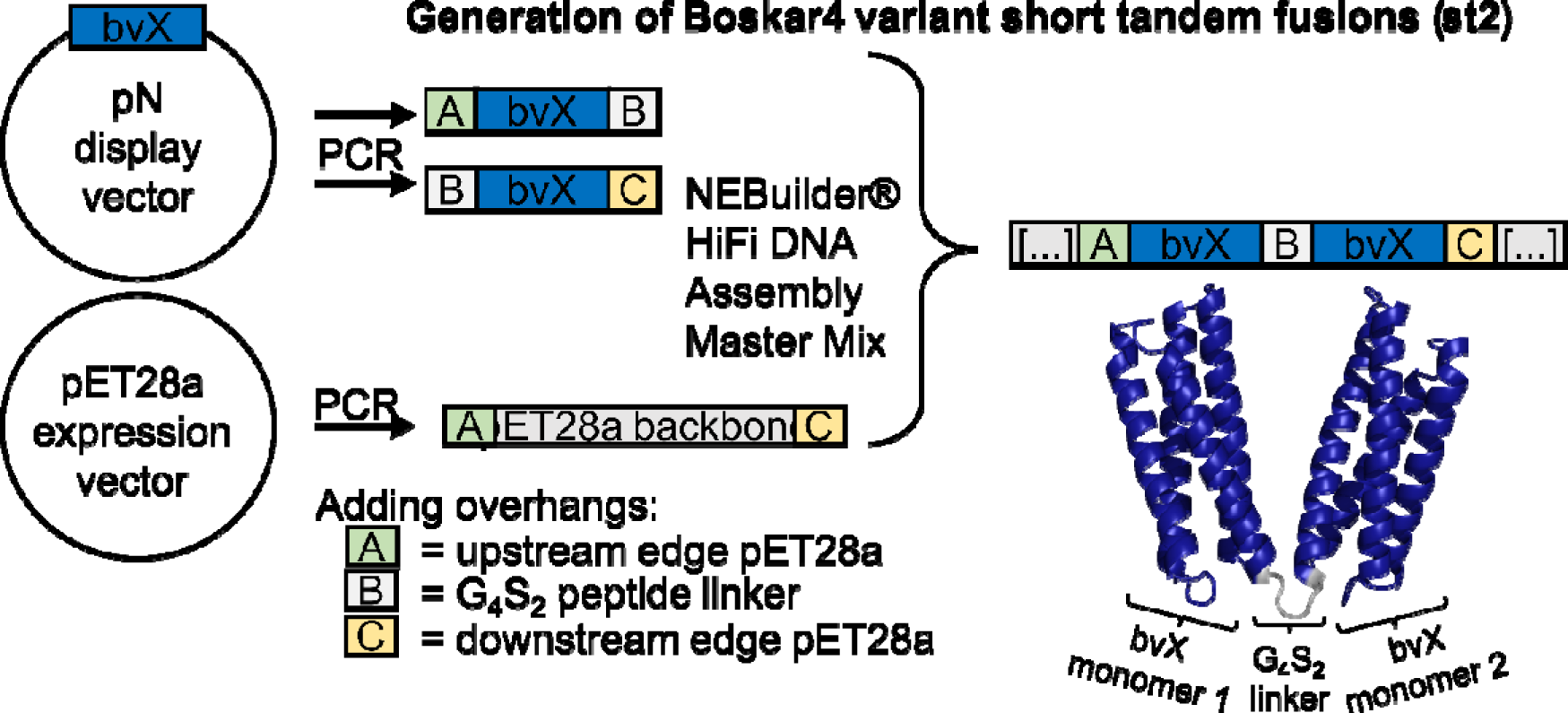
The Boskar4 variant short tandem fusions (st2) were generated by the assembly of three fragments consisting of the expression vector, two copies of a Boskar4 variant (bvX) and an overlap including the G_4_S_2_ peptide linker. The same approach was also used to generate the rigid tandem fusions (ori0 to ori4) by encoding a rigid helix linker instead of a flexible GS-peptide-linker.

**Supplementary Figure 5.**
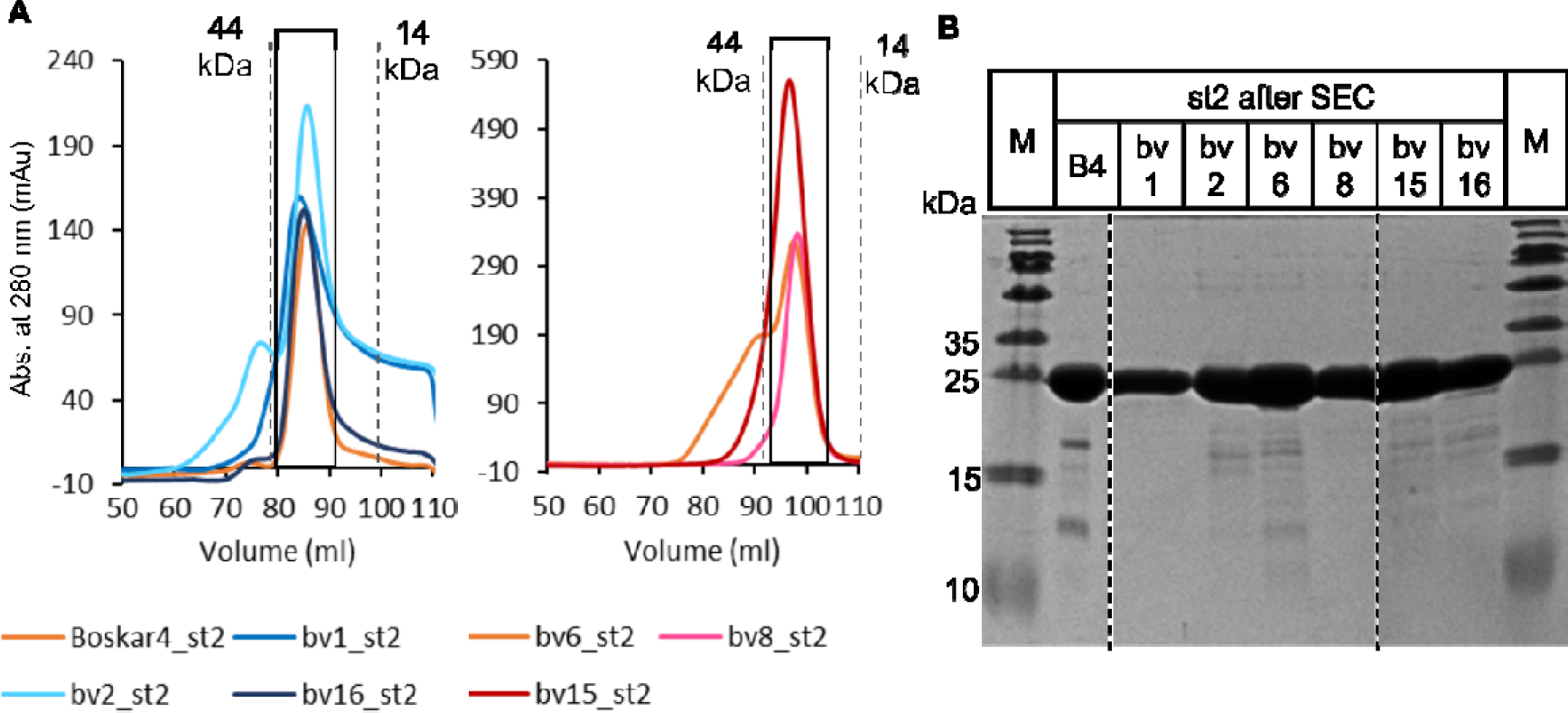
**(A)** Preparative size exclusion chromatography of Boskar4 short tandem fusion (st2) variants after Nickel IMAC on two separate SEC-columns of the same type (HiLoad 16/600, Superdex 200 pg). Dotted lines mark the molecular weight standard for the corresponding column. All fractions of the main peak (black box) were collected and concentrated by ultrafiltration (10 kDa cut-off). **(B)** A representative sample of each protein and a protein molecular weight marker (M) (11852124, Thermo Fisher Scientific) was loaded on a SDS-PAGE (18% PA, 150 V, 1 h) which was subsequently stained with Coomassie. The expected molecular weight of Boskar4_st2 and variants (with not cleaved TEV-cleavage site and His-tag) is ∼29 kDa.

**Supplementary Figure 6.**
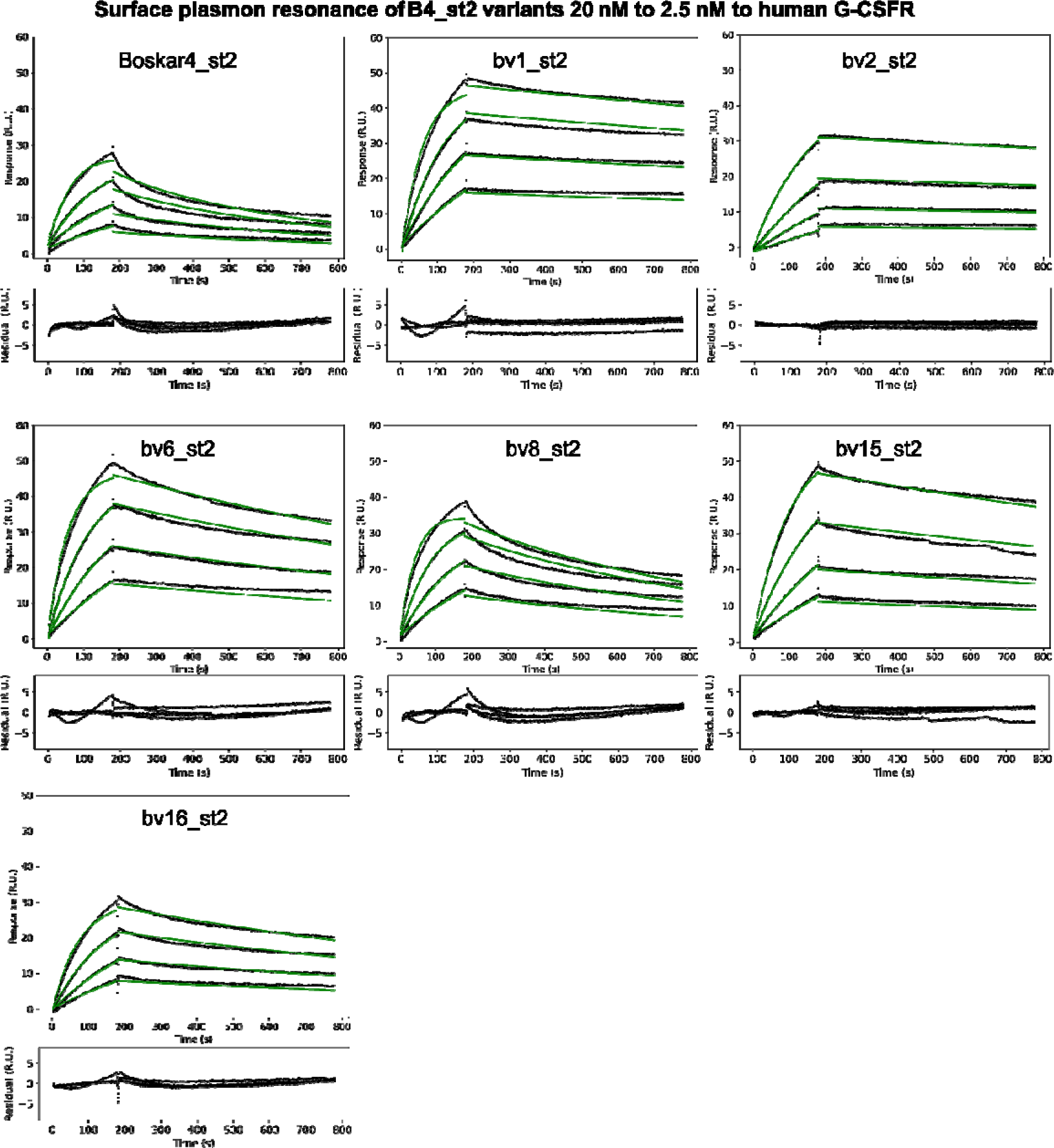
Six of the enhanced variants and Boskar4 itself (B4) were chosen to generate short tandem fusions (st2) and measured with surface plasmon resonance (SPR) against immobilized hG-CSFR. The association rate constant (*k_a_*), the dissociation rate constant (*k_d_*), and the apparent equilibrium dissociation constant (*K_D_*) were estimated from the corresponding fits (green lines) of a two-fold titration series from 20 nM to 2.5 nM (compare Table 1).

**Supplementary Figure 7.**
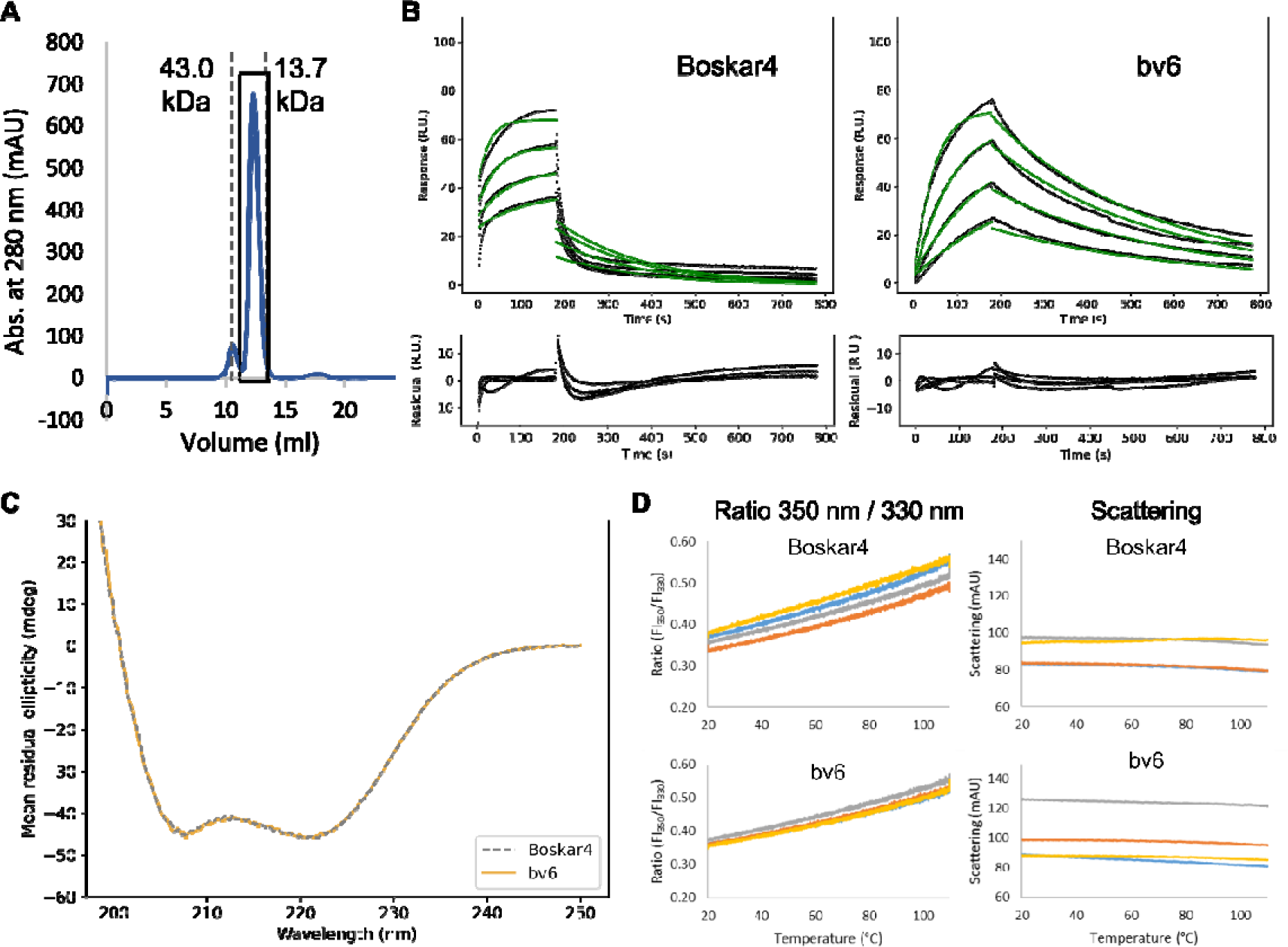
**(A)** Preparative size exclusion chromatography of monomeric bv6 on a Superdex® 75 10/300 GL with an expected size of ∼15 kDa. Dotted lines mark the molecular weight standard for the corresponding column. All fractions of the main peak (black box) were collected and concentrated by ultrafiltration (10 kDa cut-off). **(B)** Surface plasmon resonance measurement (black line) and the corresponding fit (green lines) of a two-fold dilution series of monomeric Boskar4 and the variant bv6. The highest concentration used for Boskar4 was 1250 nM and for bv6 125 nM. The obtained binding parameters are listed in Table 1. **(C)** The single-domain Boskar4 (B4; gray) and an example variant (bv6; orange) were analyzed with circular dichroism and show strong helical signal. **(D)** Thermostability measurement with nanoDSF of 1 mg/mL of monomeric Boskar4 and bv6 with 4 technical replicates, each indicating the proteins to be stable to at least 110 °C.

**Supplementary Figure 8.**
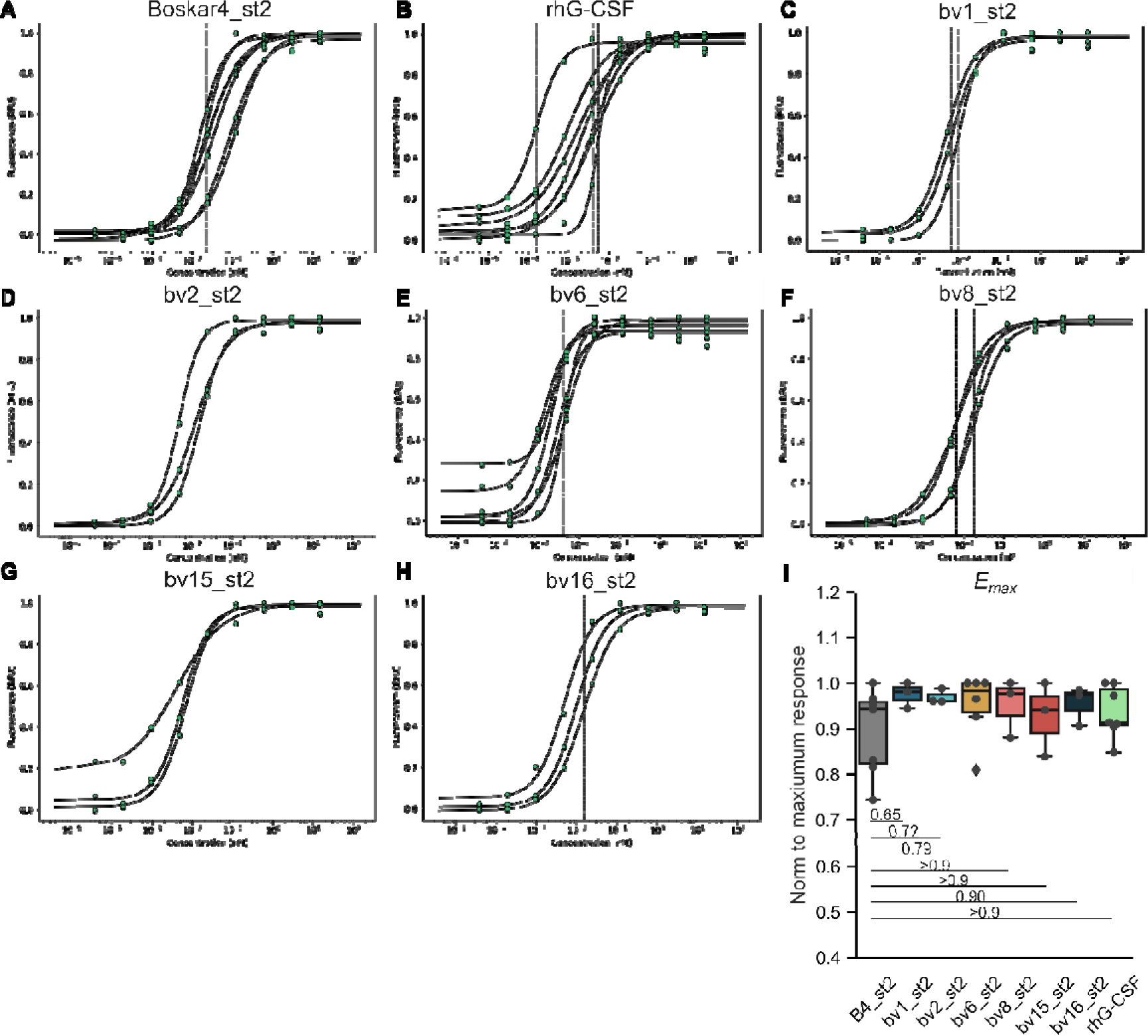
**(A-H)** Dose-response NFS-60 cell proliferation assays to determine the half-maximal effective concentration (*EC_50_*) of Boskar4_st2 variants and human recombinant G-CSF (hrG-CSF). The fits to determine the *EC_50_* (pM) of at least three independent experiments per G-CSFR agonist are shown (compare also Table 1 and Fig. 2C). **(I)** Maximum proliferative activity (*E_max_*) normalized to the maximum response within each independent experiment represented in (A-H). For statistical analysis, an ordinary one-way ANOVA was performed, followed by a Tukey HSD test.

**Supplementary Figure 9.**
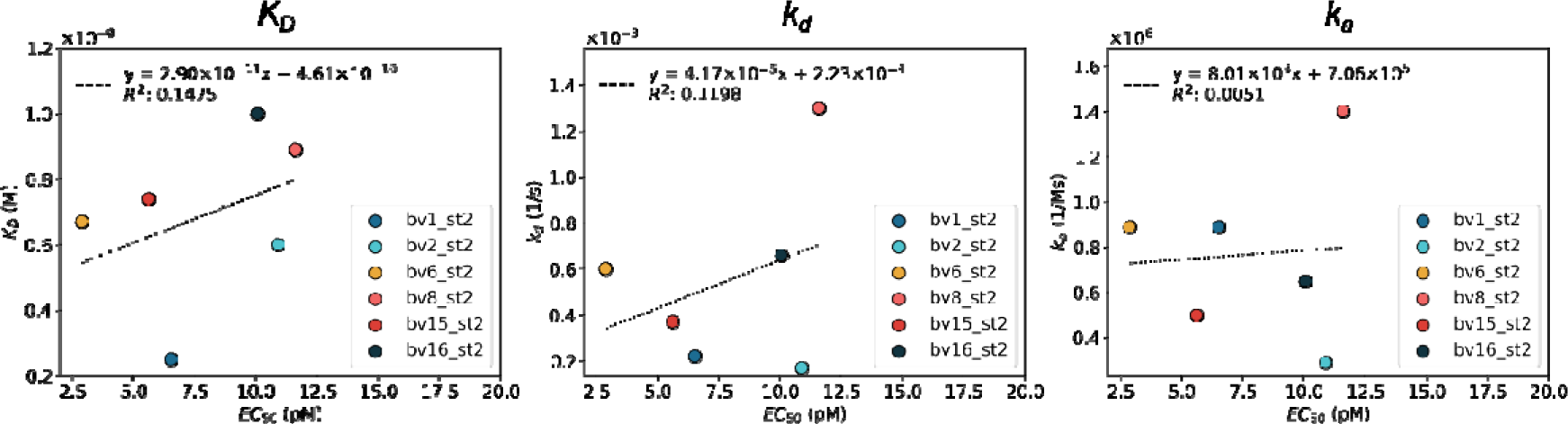
Shown is the correlation of a linear fit between the apparent affinity (*K_D_*), association rate constant (*k_a_*) or dissociation rate constant (*k_d_*) and the obtained NFS-60 activity (EC_50_) for short tandem fusions of Boskar4 (B4) or the affinity-enhanced variants (bv1, bv2, bv6, bv8, bv15, bv6). The coefficient of determination (*R^2^*) is shown in the left upper corner.

**Supplementary Figure 10.**
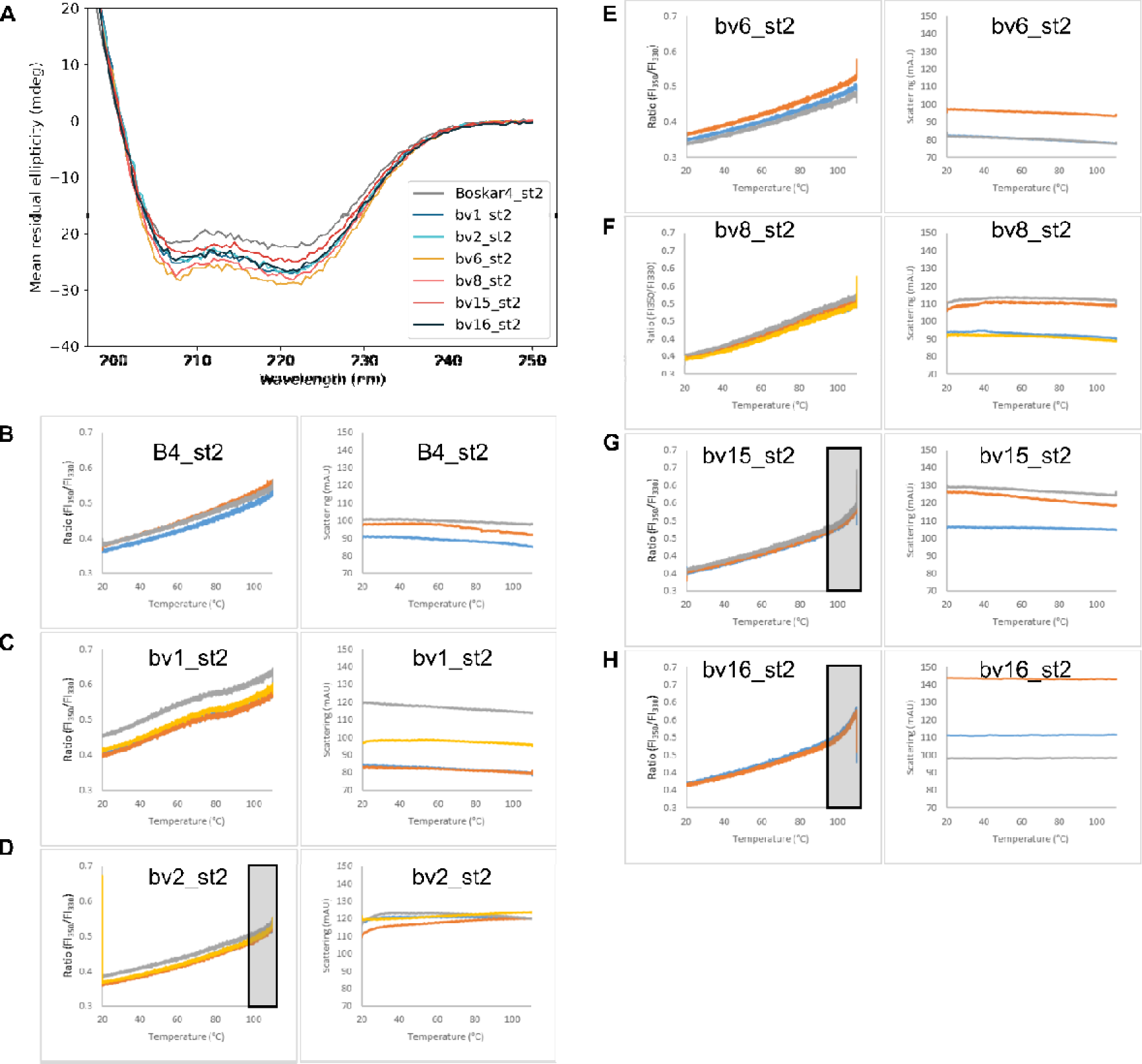
**(A)** Circular dichroism (CD) measurement of Boskar4_st2 and variants (bv1 to bv16) in PBS at 0.1 mg/mL shows expected alpha-helical signal for all samples. **(B-H)** Thermostability measurement with nanoDSF of Boskar4_st2 (B4_st2) and variants (bv1_st2 to bv16_st2) with at least 3 technical replicates. All samples were diluted to 1 mg/mL in PBS, except for bv1_st2, which was used at a concentration 0.5 mg/ml. The gray boxes indicate areas in which a melting onset was observed.

**Supplementary Figure 11.**
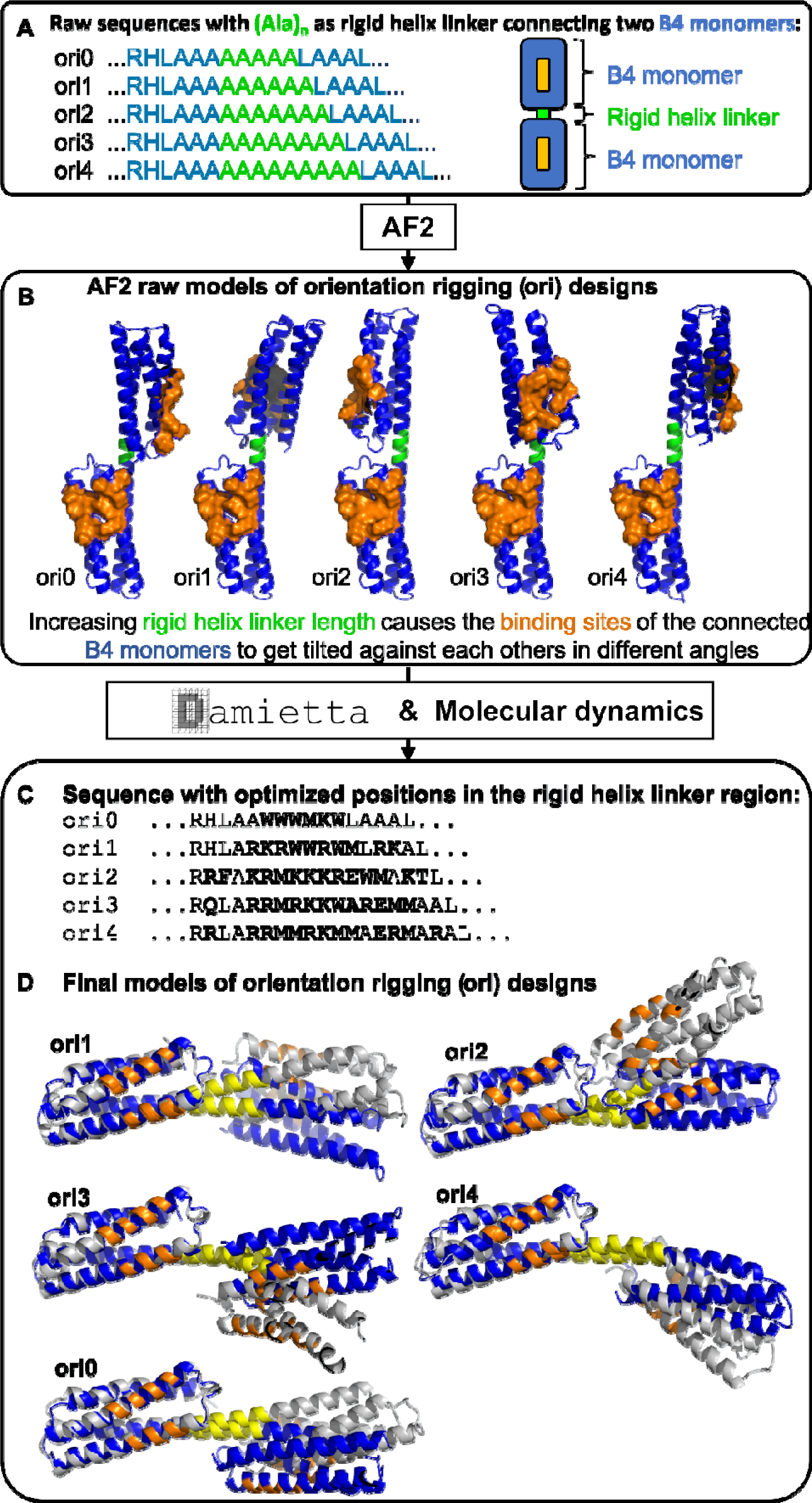
Generation of the orientation-rigging designs (oris) tuning G-CSFR activity. **(A)** Raw sequence design. **(B)** AlphaFold2 raw models. **(C)** Selected residues (bold) in the rigid helix linker region were optimized with Damietta, and the most stable designs were identified with molecular dynamics (MD). **(D)** The MD input structure of each top hit is displayed in blue, and the last frame of the MD-simulation is indicated in gray. The linker helix area is depicted in yellow, and the G-CSFR binding sites are highlighted in orange.

**Supplementary Figure 12.**
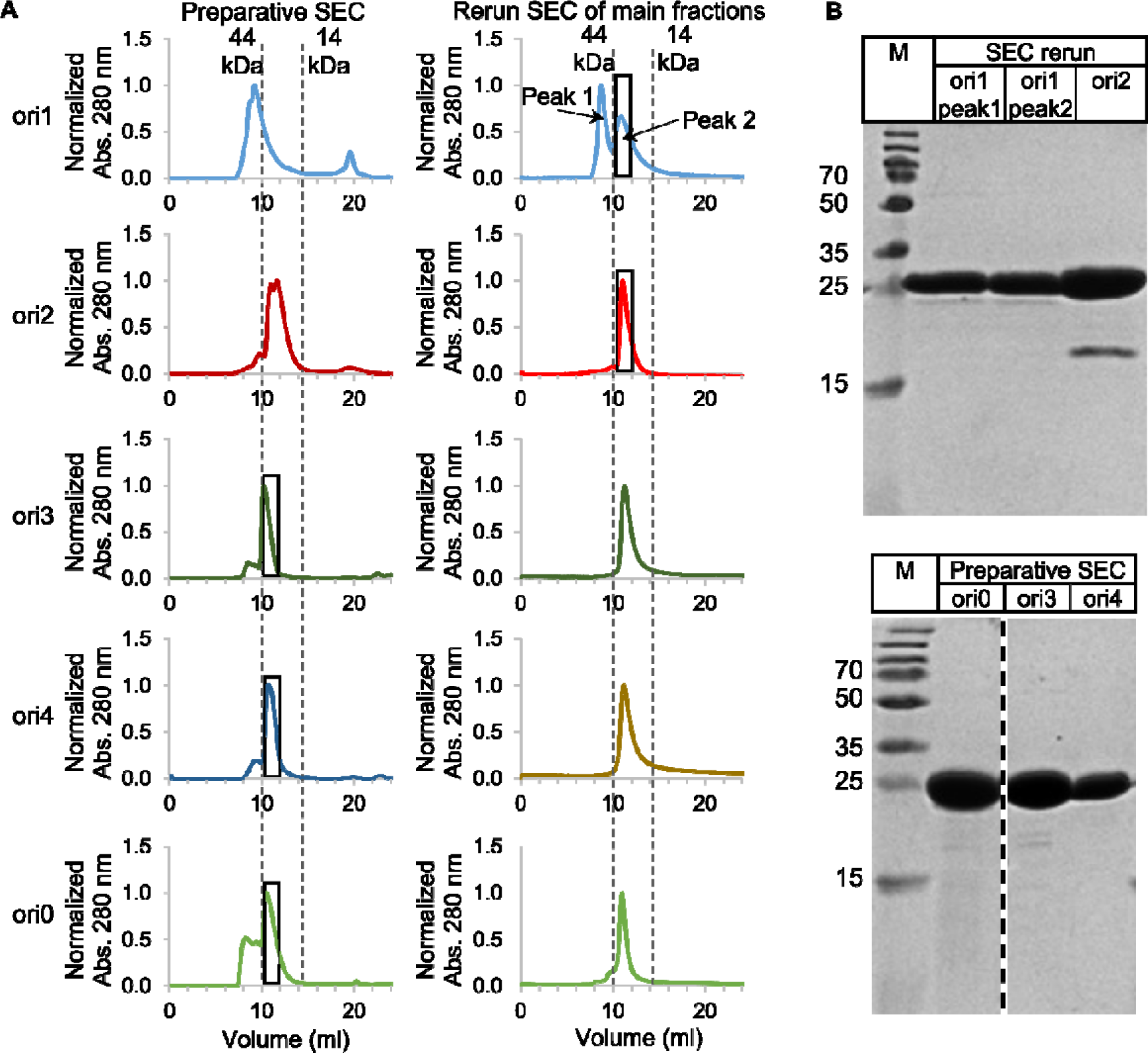
**(A)** Size exclusion chromatography of ori0, ori1, ori2, ori3 and ori4 after Nickel IMAC was performed on a Superdex Increase 75 10/300 GL (Cytiva). The dotted lines mark the molecular weight standard for the corresponding column. All fractions of the main peak were collected and concentrated by ultrafiltration (10 kDa cut-off). Part of the collected fractions were rerun on the same column to validate the purity and correct molecular mass. **(B)** A representative sample of each fraction containing the protein as used for the subsequent experiments (black boxes in (A)) was loaded on a SDS-PAGE. The expected molecular weight of the oris (with uncleaved TEV-cleavage site and His-tag) is ∼29 kDa. For ori2, the main fraction was collected after the SEC rerun and for ori1, peak 1 and peak 2 were collected separately. The main fractions after the preparative size exclusion of ori0, ori3 and ori4 were directly used for subsequent experiments.

**Supplementary Figure 13:**
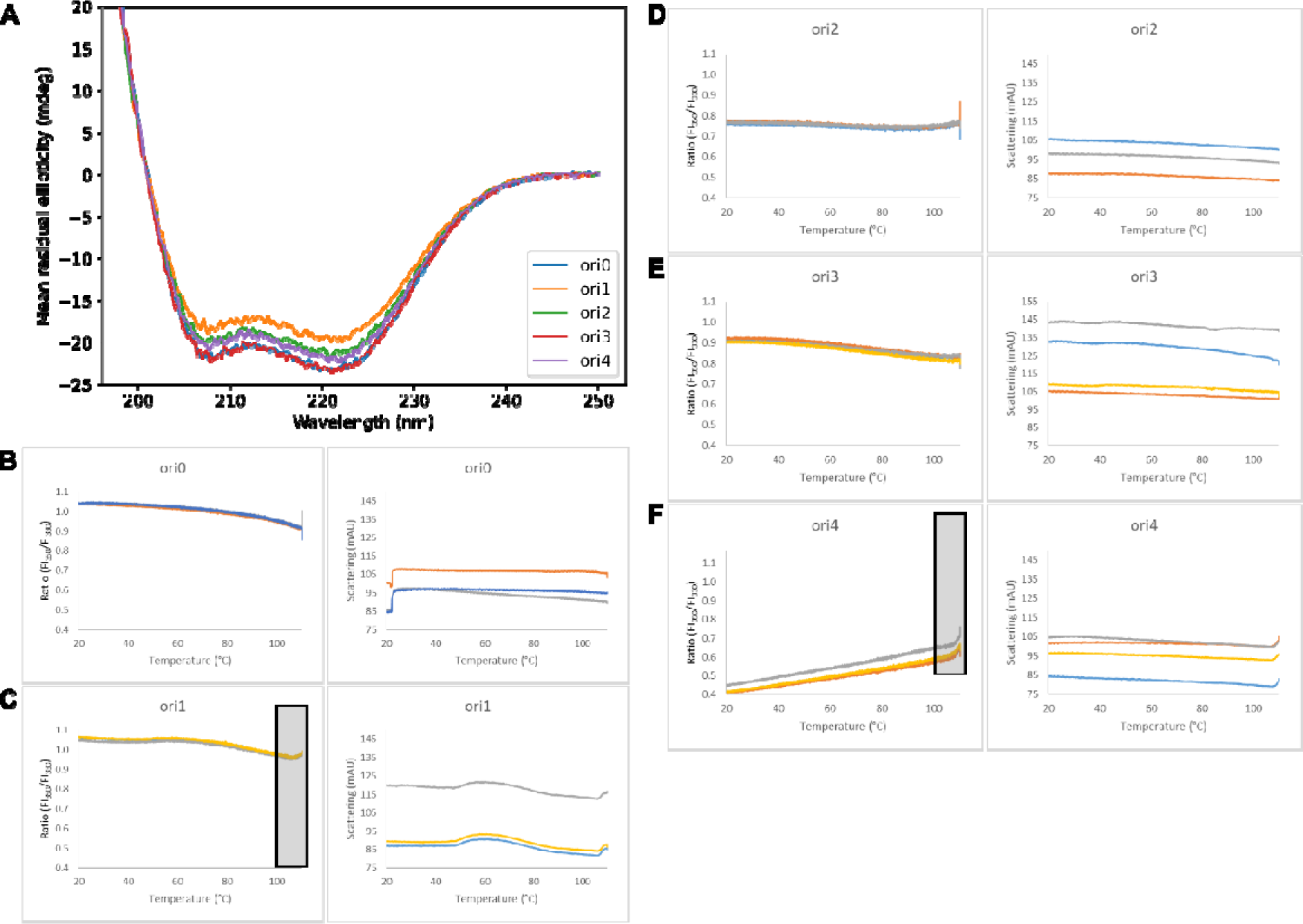
**(A)** Circular dichroism (CD) measurement of ori0, ori1, ori2, ori3 and ori4 in PBS at 0.1 mg/mL showed the expected alpha-helical signal for all samples. **(B-F)** Thermostability measurement with nanoDSF of the rigid tandem fusions (ori0 to ori4) with at least 3 technical replicates. All samples were diluted to 1 mg/mL in PBS. The gray boxes indicate areas in which a melting onset was observed.

**Supplementary Figure 14.**
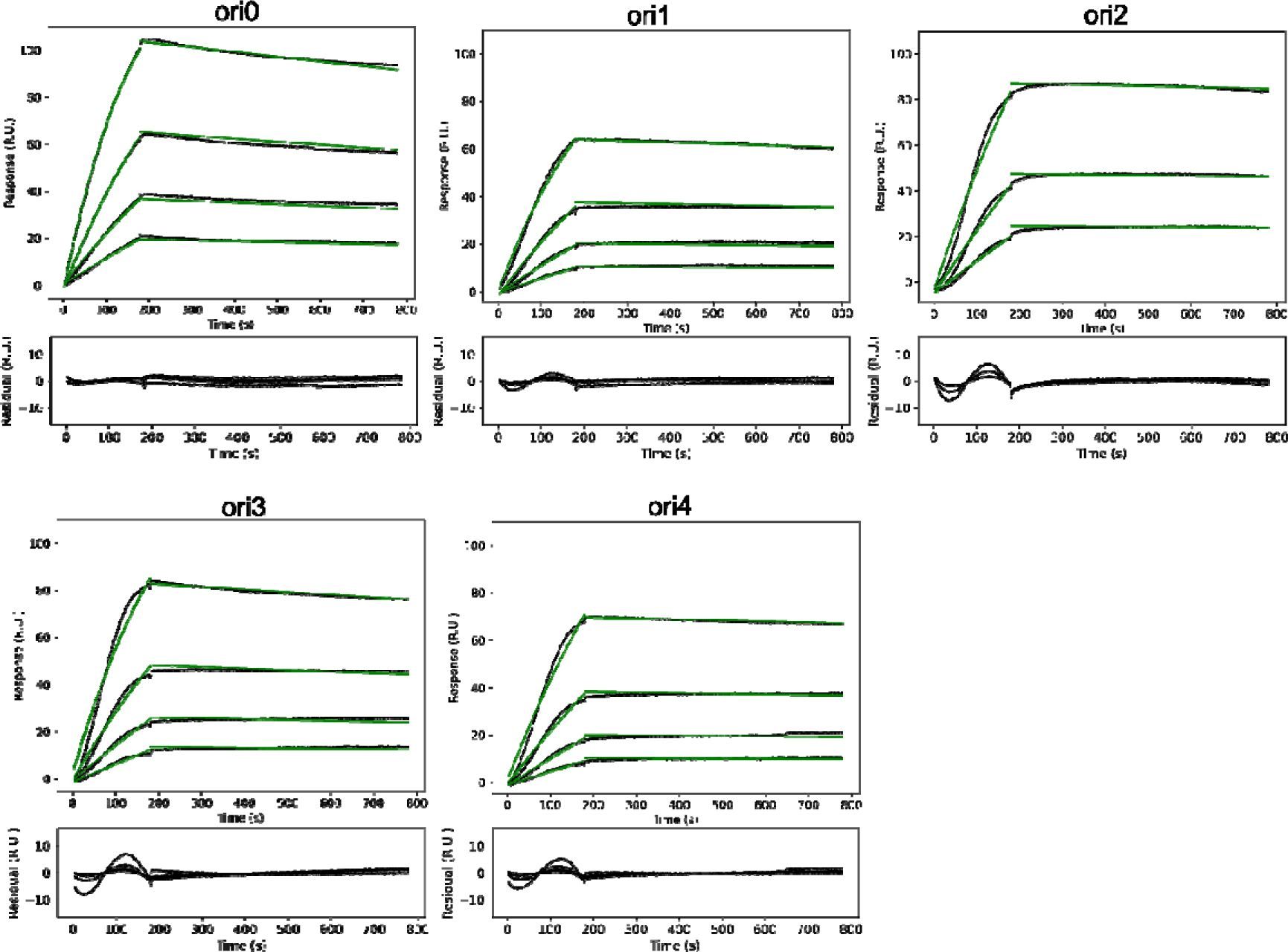
Surface plasmon resonance (SPR) measurement of oris against immobilized hG-CSFR. The association rate constant (*k_a_*), the dissociation rate constant (*k_d_*), and the apparent equilibrium dissociation constant (*K_D_*) were estimated from the corresponding fits (green lines) of a titration series ranging from 20 nM to 2.5 nM (compare Table 1).

**Supplementary Figure 15.**
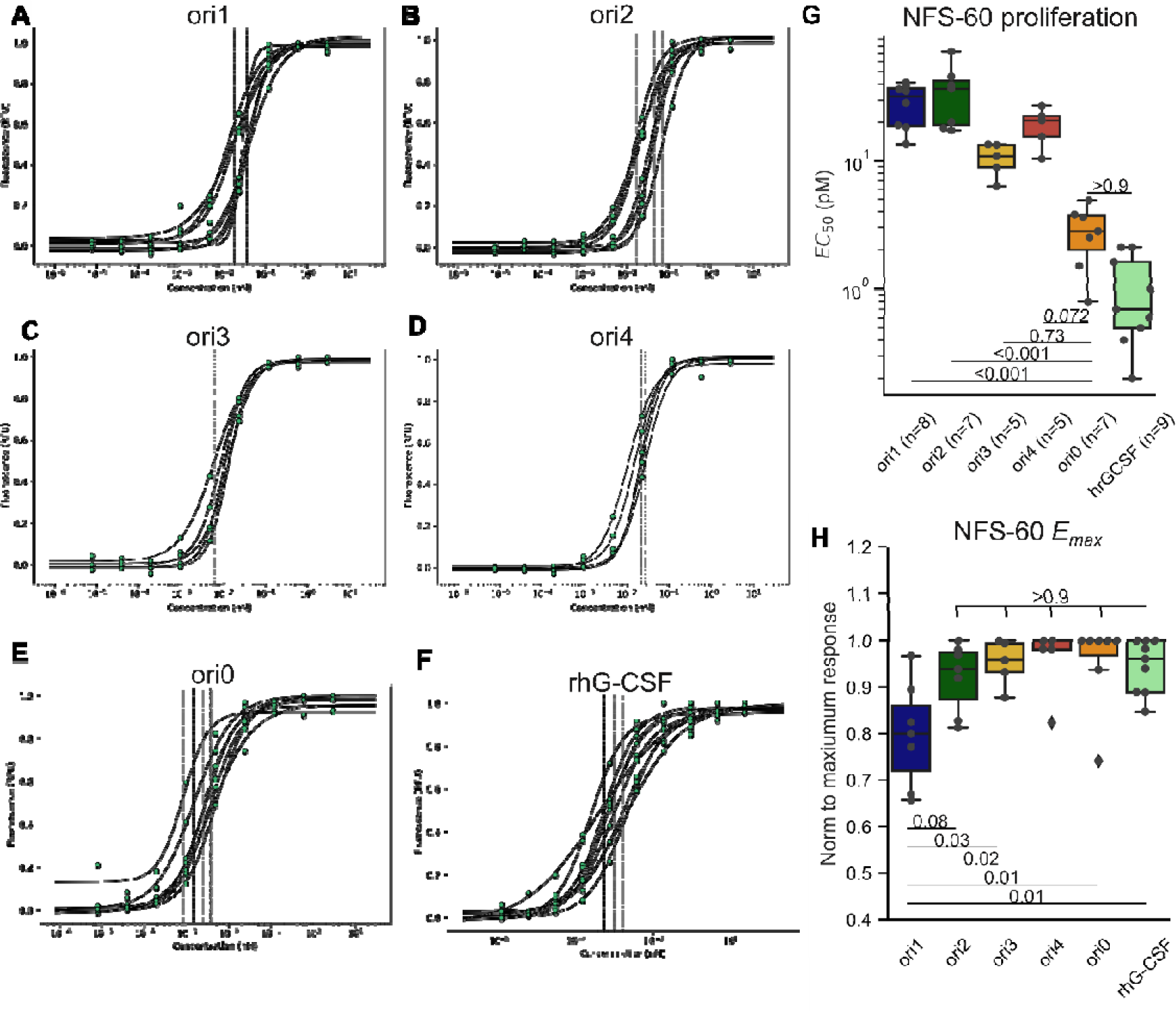
**(A-F)** Dose-response NFS-60 cell proliferation assays to determine the half-maximal effective concentration (EC_50_) of the oris and G-CSF (rhG-CSF, Lenograstim). Given are the fits to determine the *EC_50_* of at least five independent experiments per G-CSFR agonist (compare also Table 1 and Fig. 3C). **(G, H)** Box plot of the *EC_50_* (pM) values (G) and the maximum proliferative activity (*E_max_*) normalized to the maximum response (H) within each independent experiment represented in (A-F). For statistical analysis in (G) and (H), ordinary one-way ANOVA, followed by a Tukey HSD test was performed.

**Supplementary Figure 16.**
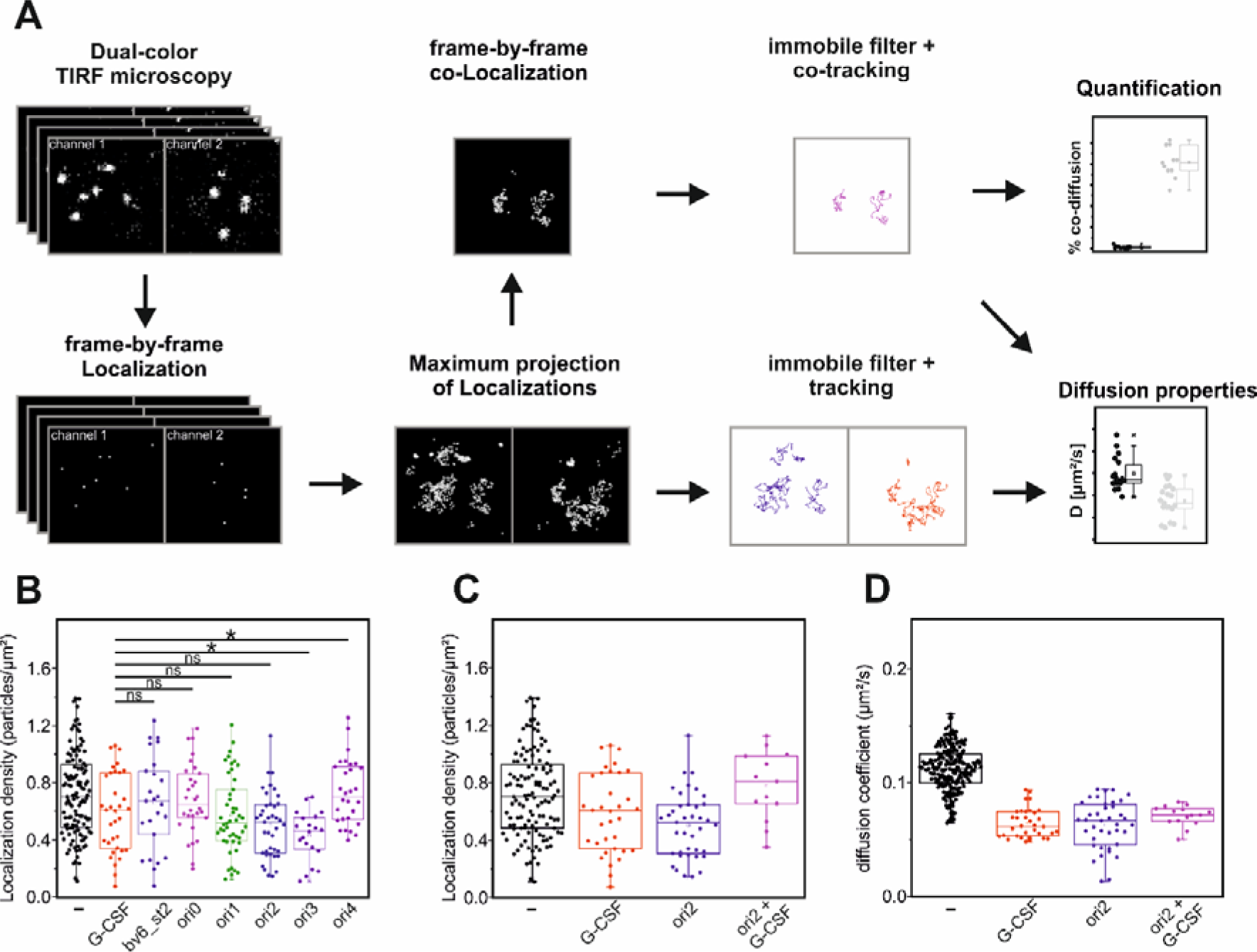
The dimerization of G-CSFR was analyzed with single molecule imaging for G-CSF and the design agonists. **(A)** Schematic representation of Single molecule co-tracking (SMCT) workflow. **(B,C)** Receptor density distribution for SMCT experiments. **(D)** Diffusion coefficients of G-CSFR and induced complexes in competition assay.

**Supplementary Figure 17.**
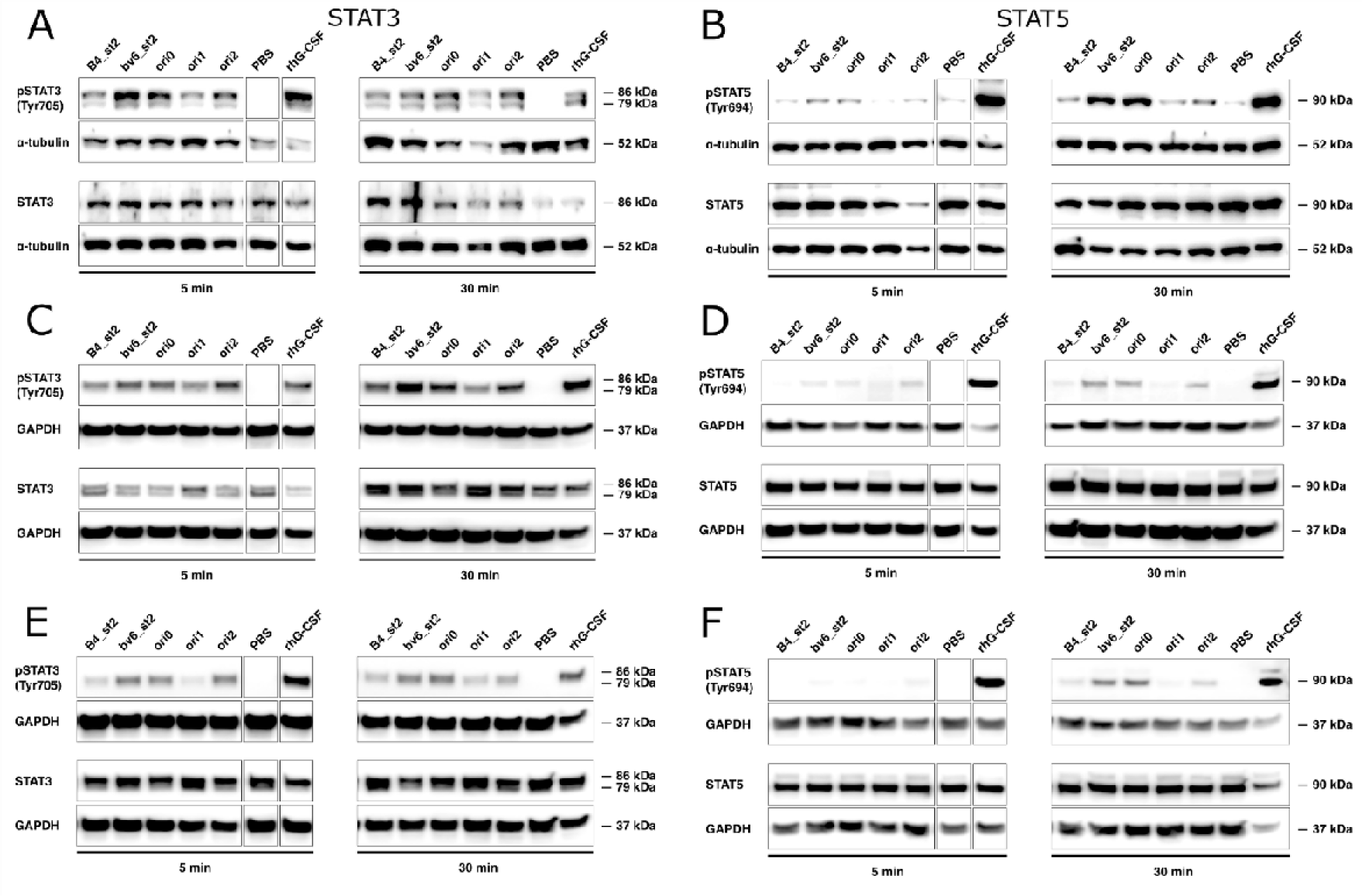
**(A-F)** Western Blot analysis of extracts from NFS-60 cells treated with B4_st2, bv6_st2, ori0, ori1, ori2 designs, PBS, or rhG-CSF for 5 or 30 min, using phospho-STAT3 (Tyr705) (A, C, E) or phospho-STAT5 (Tyr694) (B, C, D) rabbit antibodies. As loading control, α-tubulin rabbit antibody or GAPDH rabbit antibody was used. A and B, C and D, E and F represent independent experiments.

**Supplementary Figure 18.**
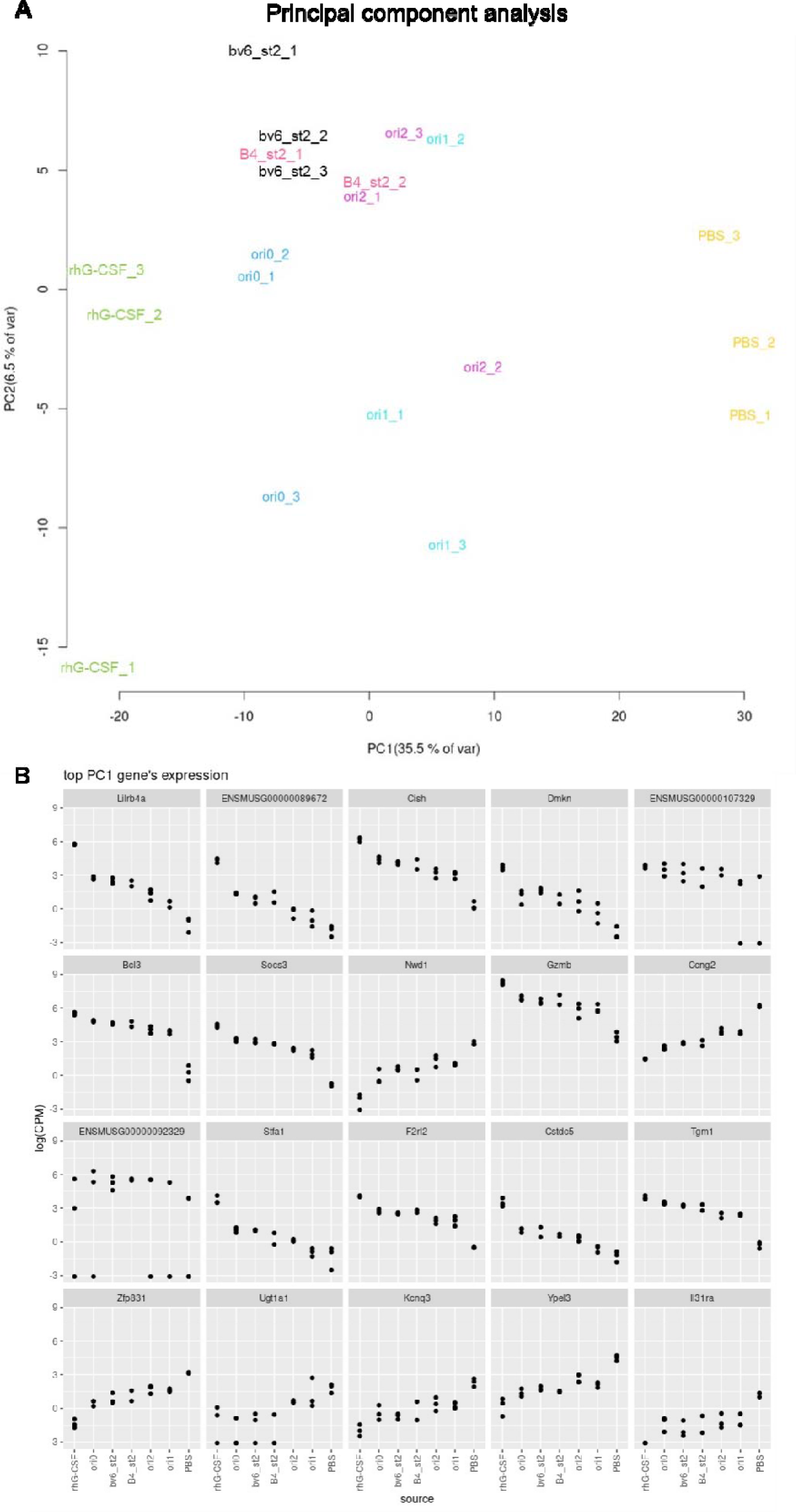
**(A)** Principal component analysis (PCA) of groups treated with rhG-CSF, agonist designs (B4_st2, bv6_st2, ori0, ori1, ori2) or PBS and **(B)** the top 20 differentially expressed genes in PC1 of PCA.

**Supplementary Figure 19.**
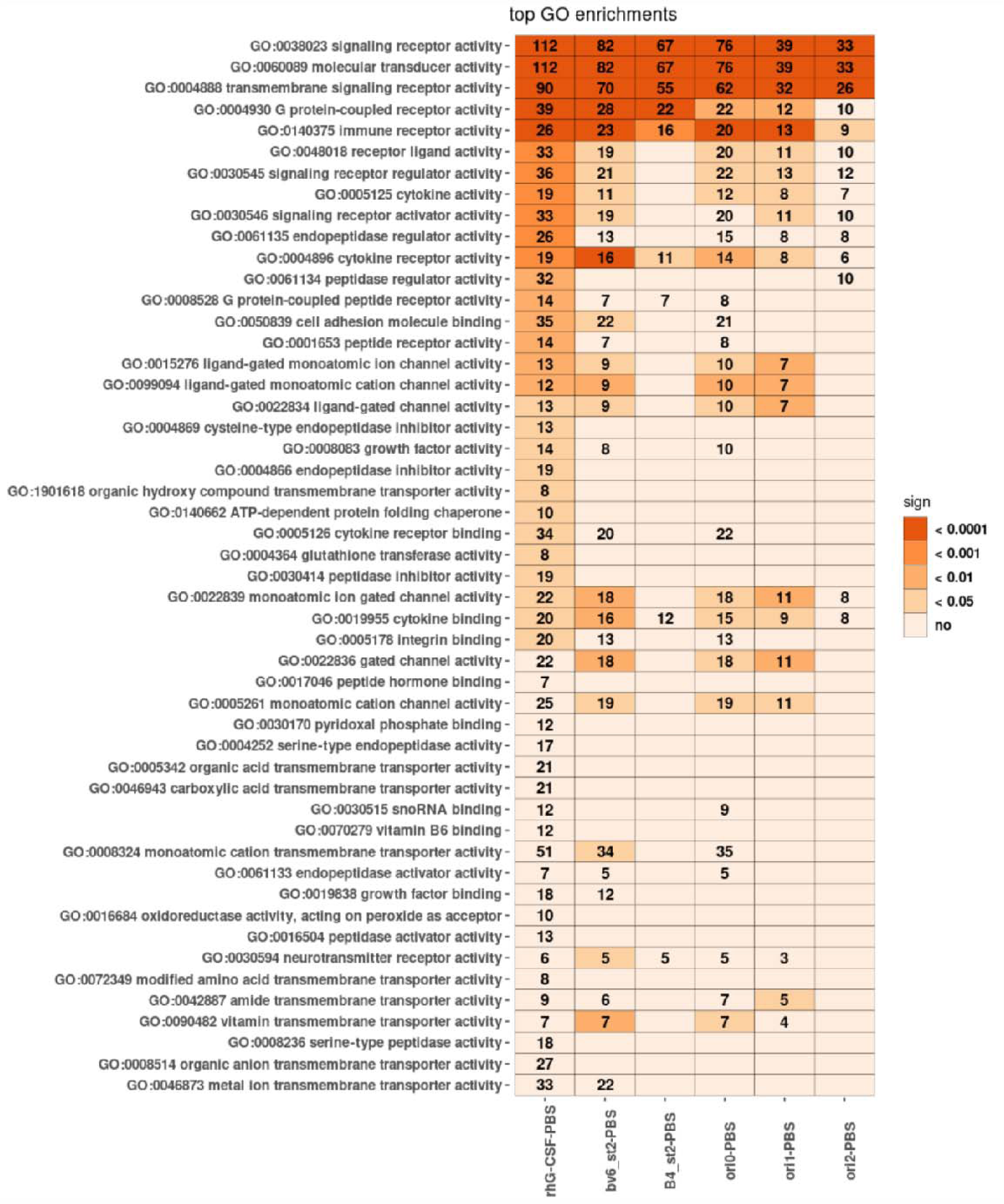
Top 50 GO pathways in the groups treated with rhG-CSF or agonist designs (B4_st2, bv6_st2, ori0, ori1, ori2) compared to PBS.

**Supplementary Figure 20.**
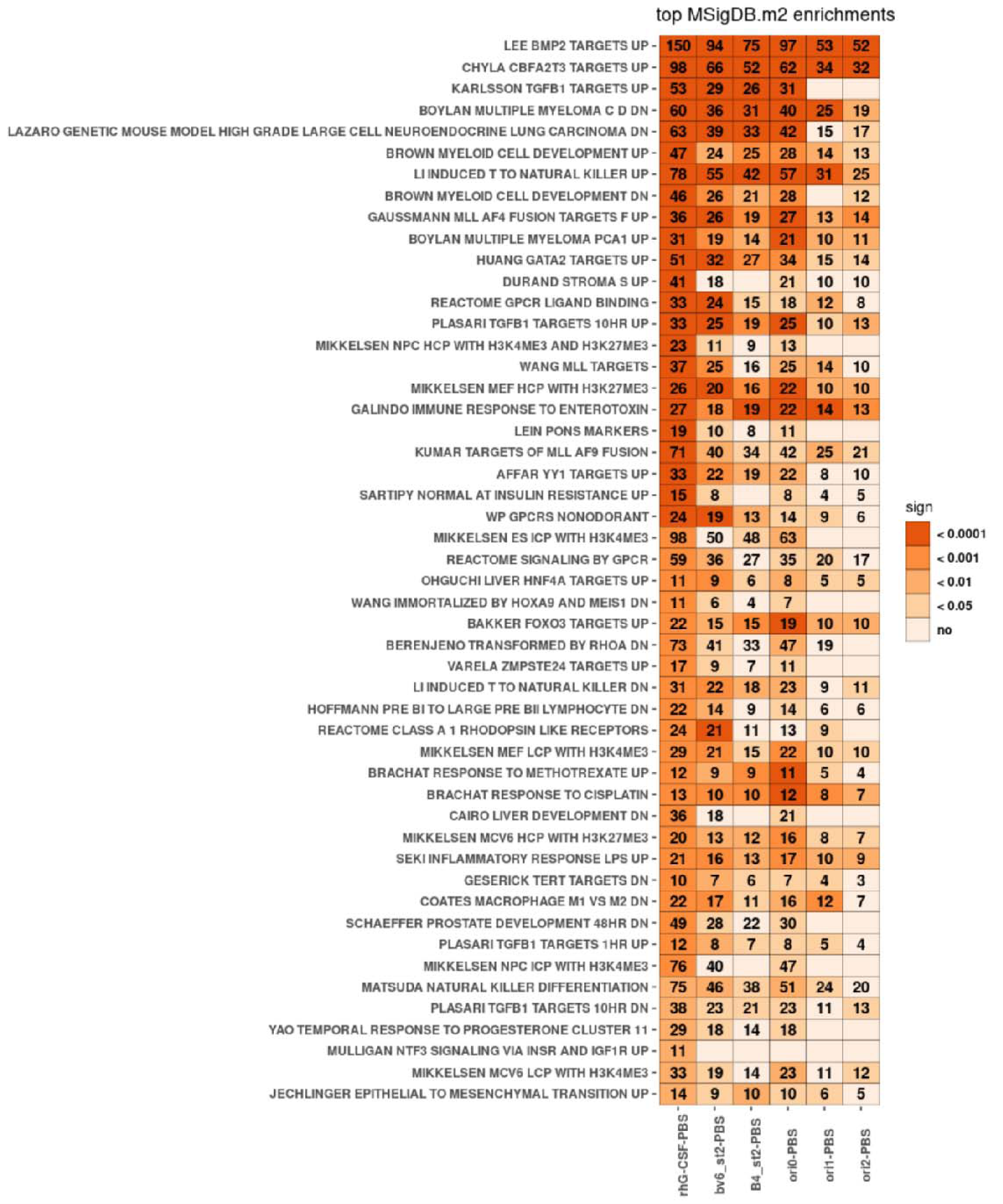
Top 50 MSigDB.m2 pathways in the groups treated with rhG-CSF or agonist designs (B4_st2, bv6_st2, ori0, ori1, ori2) compared to PBS.

## Supplementary tables

**Supplementary Table 2.**
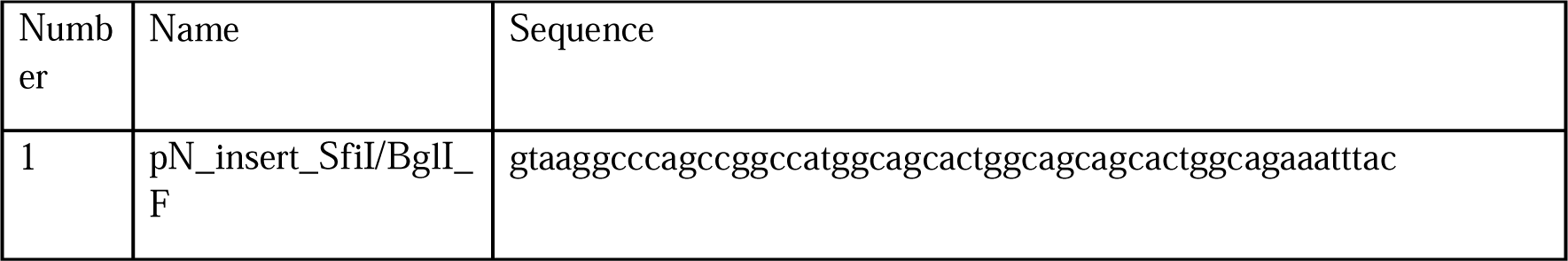
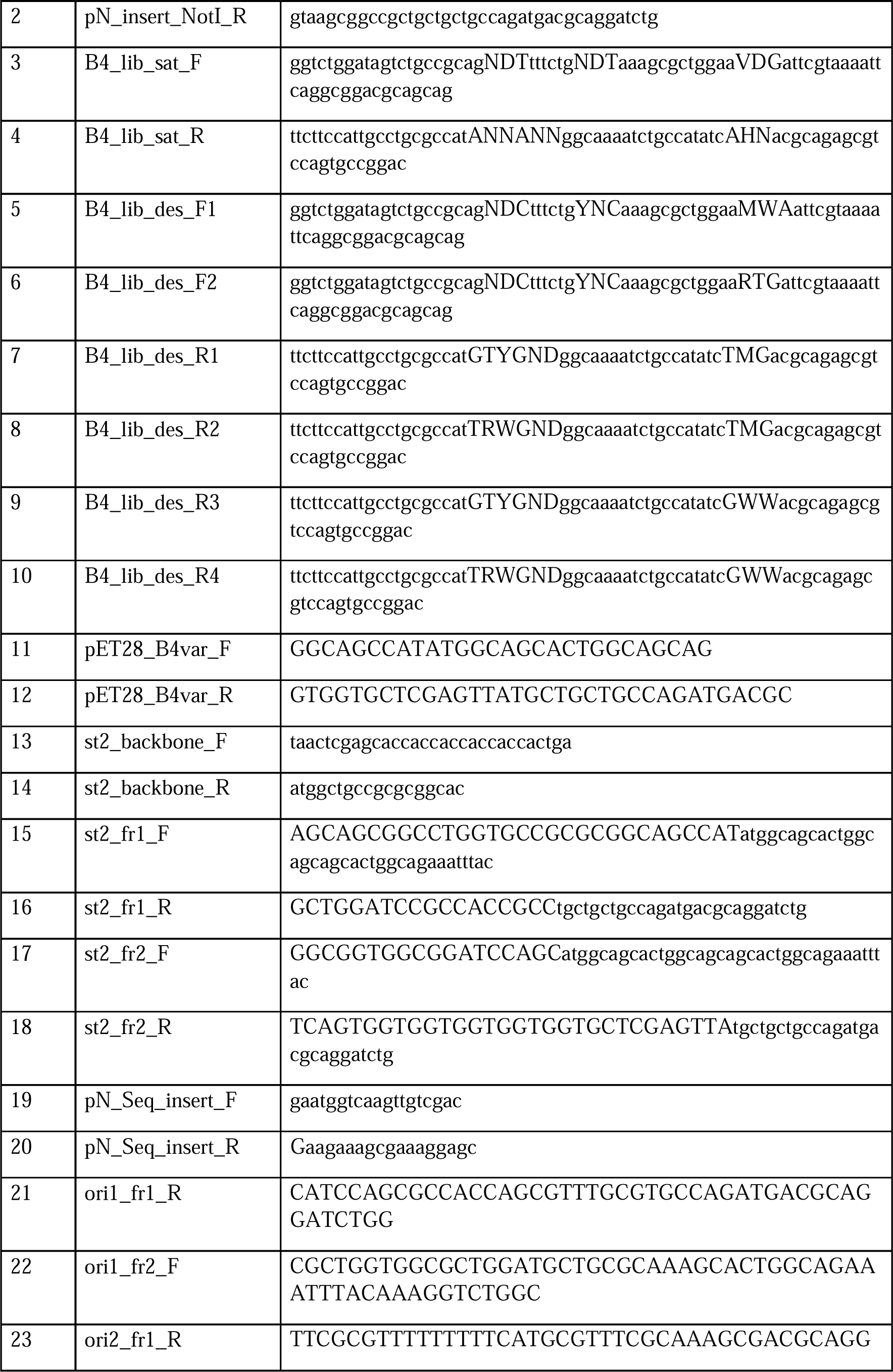
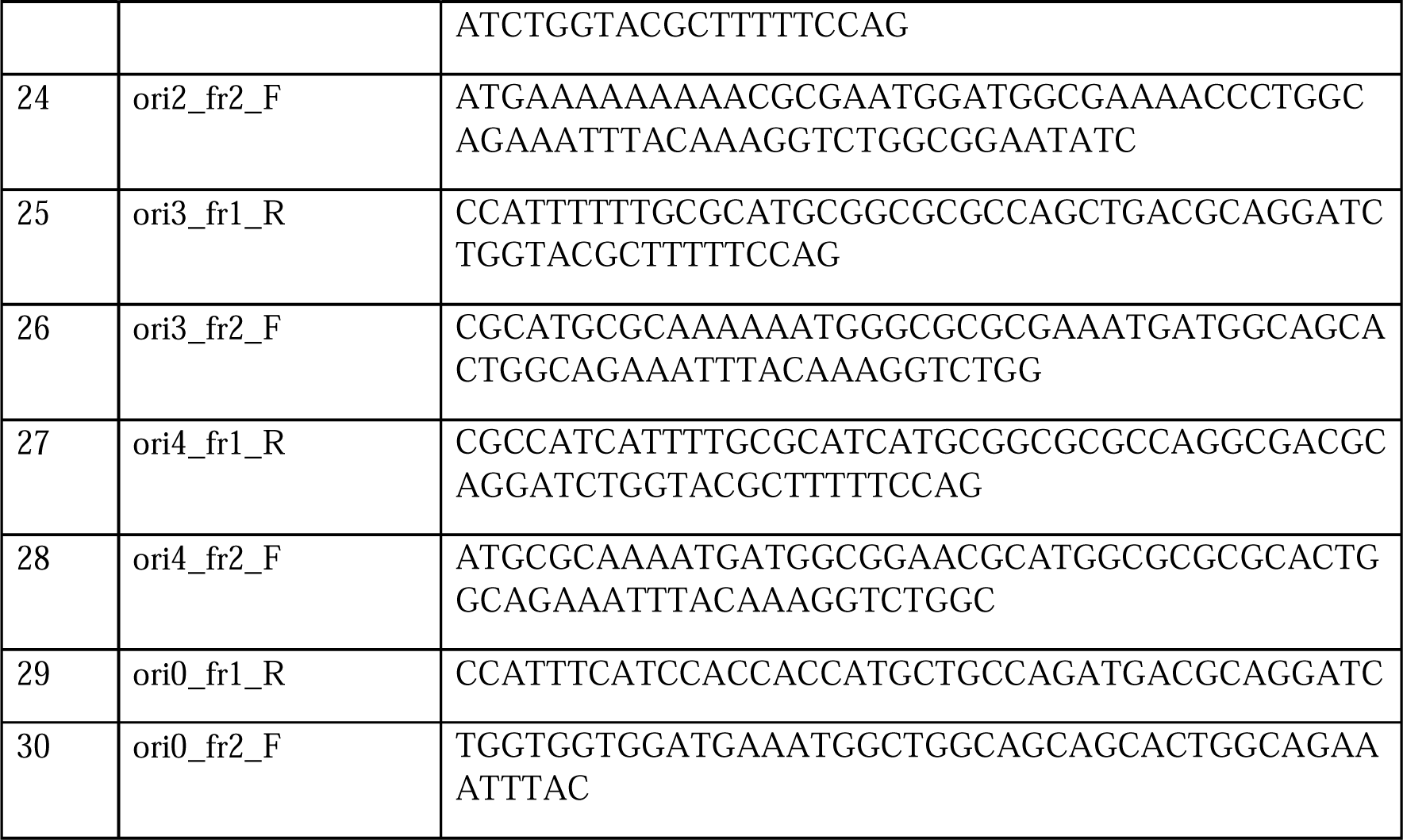
Primers used in this study.

**Supplementary Table 3:** Composition of PCR reaction mix used for all reactions with the following specifications; for all reactions 5 ng/µL template was inputted but for the ones whose products were used for assembly reactions, 0.3 ng/mL were inputted. In the case of the use of multiple primers for the library backbone PCR, all F or R primers of each construct were mixed at a total concentration of 10 µM (e.g. 2.5 µM of primer 7 to 10 each). PCR program was used for all reactions either with 60 seconds extension time for all insert fragments or 340 seconds for all backbone fragments.

| PCR reaction mix | | | Volume (μL) |
| --- | --- | --- | --- |
| dNTPs 10 mM each |  |  | 1 |
| Template 0.3 ng/μL or 5 ng/μL | | | 1 |
| Primer F 10 μM | | | 2.5 |
| Primer R 10 μM | | | 2.5 |
| 5x Q5® Reaction Buffer (#B9027, NEB) |  |  | 10 |
| GC enhancer |  |  | 10 |
| Q5® High-Fidelity DNA Polymerase (#M0491S, NEB) |  |  | 0.5 |
| H<sub>2</sub>O ad 50 μl | | | 22.5 |
| PCR program | Temperature (°C) | Time (s) | Cycle |
| Initial denaturation | 98 | 30 | 1x |
| Denaturation | 98 | 10 | 30x |
| Extension | 72 | 60 or 340 |  |
| Final Extension | 72 | 300 | 1x |
| Hold | 8 | - | 1x |

**Supplementary Table 4:**
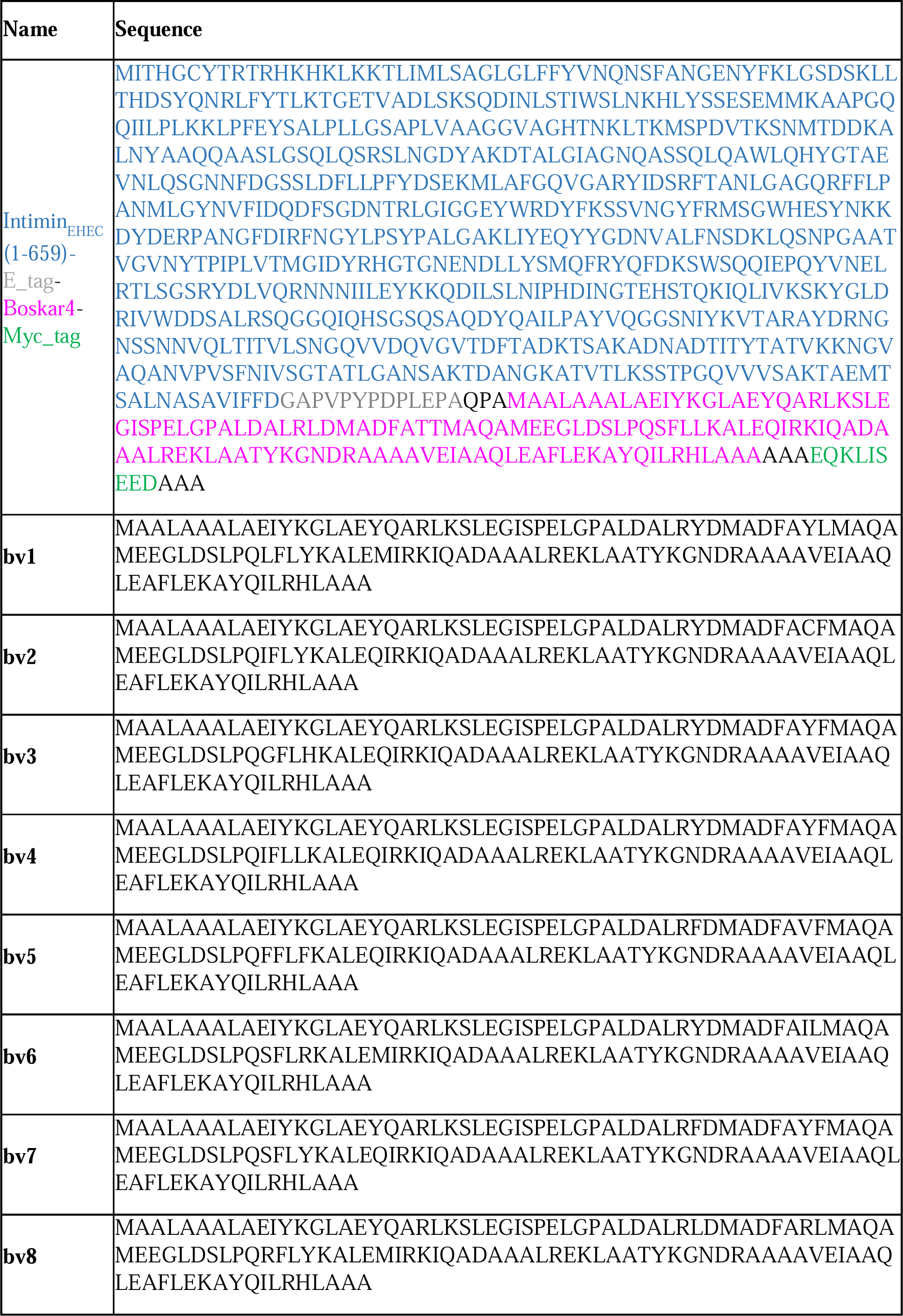
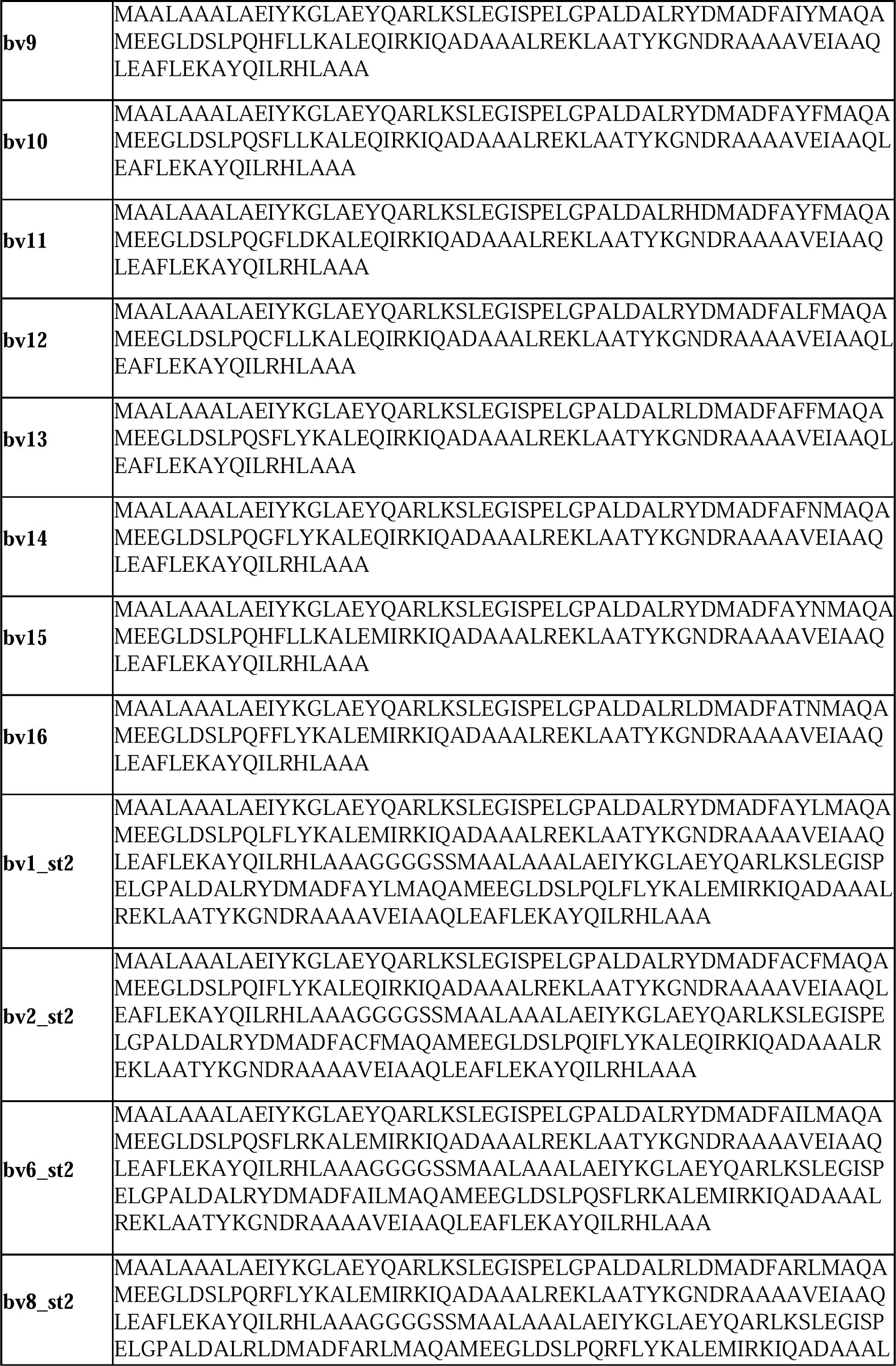
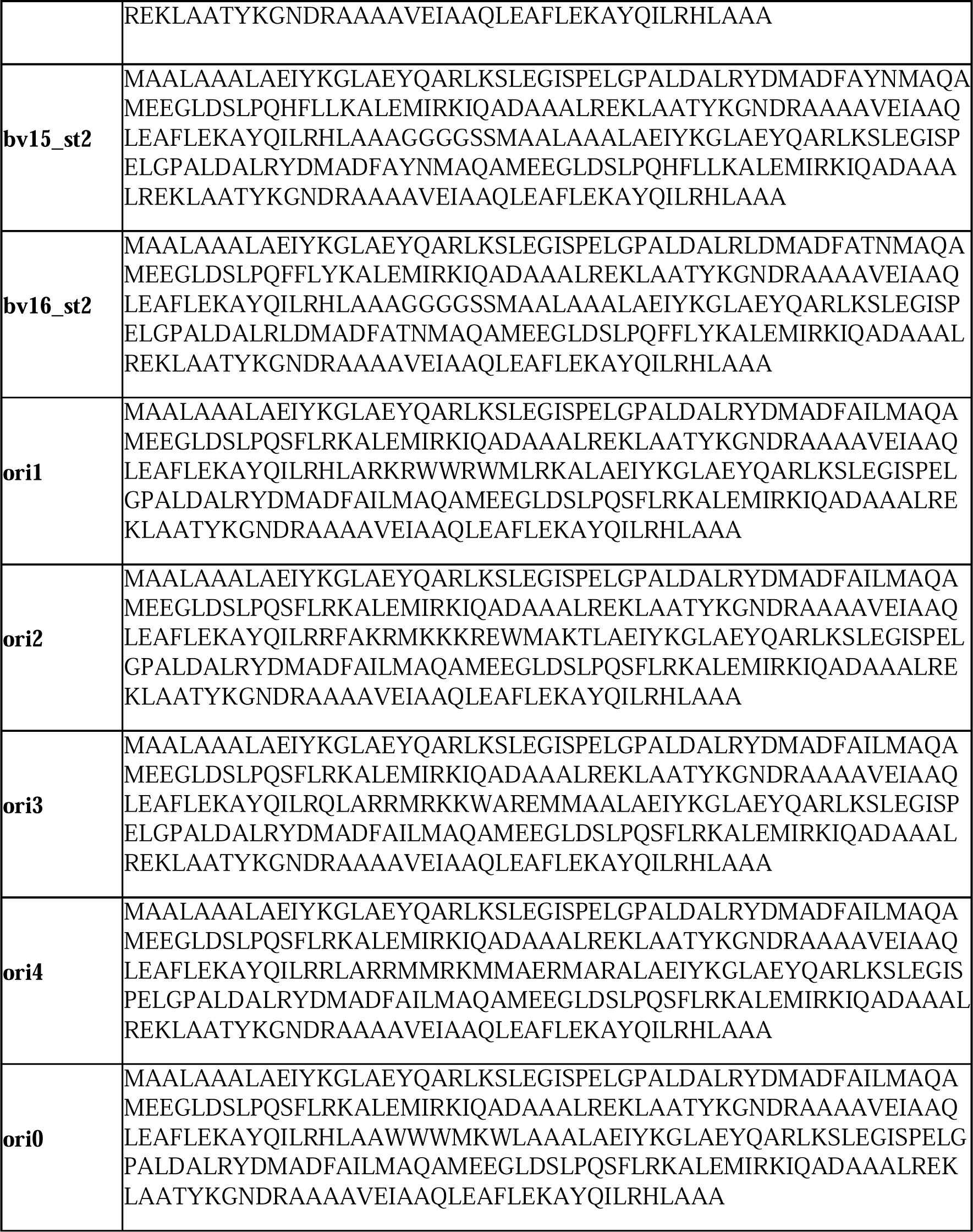
Sequence of the fusion protein between Intimin [35] and Boskar4 coded by pNB4 (colored) and sequences of all Boskar4 variants and designs (black) described in this study

## Supplementary protocols

### Damietta specifications for the design of orientation-rigging ligands

Example Damietta specifications for weights, sample parameters, the mutable residues, and repackable residues for designing the rigid linker helix region of ori0 (compare Material and Methods, Design of orientation-rigging ligands):

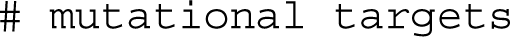
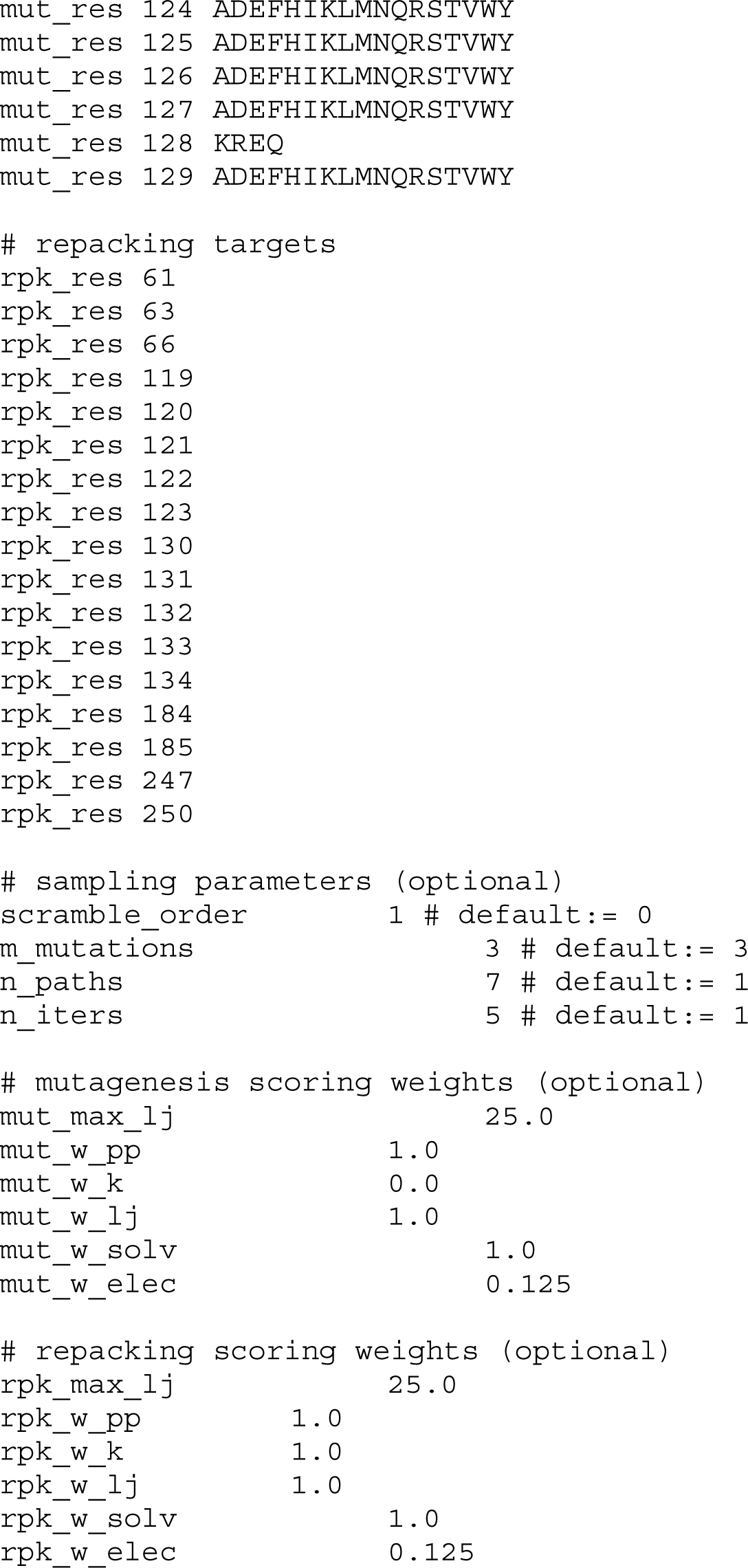

